# Regionally distinct trophoblast regulate barrier function and invasion in the human placenta

**DOI:** 10.1101/2022.03.21.485195

**Authors:** Bryan Marsh, Yan Zhou, Mirhan Kapidzic, Susan Fisher, Robert Blelloch

## Abstract

The human placenta contains two specialized regions: the villous chorion where gases and nutrients are exchanged between maternal and fetal blood, and the smooth chorion which surrounds more than 70% of the developing fetus but whose cellular composition and function is poorly understood. Here, we use single cell RNA sequencing to compare the cell types and molecular programs between these two regions in the second trimester human placenta. Each region consists of progenitor cytotrophoblasts (CTBs) and extravillous trophoblasts (EVTs) with similar gene expression programs. While CTBs in the villous chorion differentiate into syncytiotrophoblasts, they take an alternative trajectory in the smooth chorion producing a previously unknown CTB population which we term smooth-chorion-specific CTBs (SC-CTBs). Marked by expression of region-specific cytokeratins, the SC-CTBs form a stratified epithelium above a basal layer of progenitor CTBs. They express epidermal and metabolic transcriptional programs consistent with a primary role in defense against physical stress and pathogens. Additionally, we show that SC-CTBs closely associate with EVTs and secrete factors that inhibit the migration of the EVTs. This restriction of EVT migration is in striking contrast to the villous region where EVTs migrate away from the chorion and invade deeply into the decidua. Together, these findings greatly expand our understanding of CTB differentiation in these distinct regions of the human placenta. This knowledge has broad implications for studies of the development, functions, and diseases of the human placenta.

**Impact Statement:** Single cell RNA-sequencing of distinct regions of the human placenta identifies a smooth chorion-specific cytotrophoblast population responsible for unique functions of the smooth chorion, including acting as a barrier and restricting invasion.

## Introduction

The human placenta is the first organ to develop and forms the essential bridge between maternal and fetal tissues beginning at implantation (Knöfler et al., 2019; Turco and Moffett, 2019). The placenta must begin development rapidly upon conception in order to perform the roles of future organ systems that have not yet developed and matured in the fetus, including nutrient and oxygen transport and protection from mechanical and pathogenic insults. The placenta also performs unique functions such as modulation of maternal tolerance and hormone production (Maltepe and Fisher, 2015; Knöfler et al., 2019; Turco and Moffett, 2019). Placental development begins with the generation of stem villi surrounding the entire embryo and then proceeds asymmetrically to produce two distinct regions. At the human implantation site, the embryonic pole, the villi grow and branch to give rise to the region essential for the exchange of gases and nutrients, the chorion frondosum (also known as the chorionic villi or the villous chorion [VC]). This region includes the placental villi and the invasive EVTs. The villi located on the opposite side, the abembryonic pole, degenerate resulting in a smooth surface lacking villi termed the chorion leave (also known as the smooth chorion [SC]). The smooth chorion fuses with the amnion forming the chorionamniontic membranes (also known as the fetal membranes)[Hamilton and Boyd, 1960; Boyd and Hamilton, 1967; Benirshcke et al., 2006].

Both the villous chorion and smooth chorion are comprised of fetal derived cytotrophoblasts (CTBs), with CTBs in the villous chorion differentiating to either multinucleate syncytiotrophoblast (STB) or to invasive extravillous trophoblast (EVT)[Knöfler et al., 2019]. Compared to the villous chorion, little effort has been made to analyze types and functions of the CTBs that comprise the smooth chorion (Benirshcke et al., 2006, Garrido-Gomez et al., 2017). The CTBs of the smooth chorion exist in an epithelial like structure and lack STBs and proximity to the fetal vasculature, and thus cannot function in a manner comparable to villi in villous chorion (Benirschke et al., 2006). Furthermore, in contrast to the villous chorion where EVTs invade and remodel the maternal arteries, the cells of the smooth chorion do not invade the adjacent decidua and the maternal blood vessels it contains (Genbacev et al., 2015). Thus, the function of these smooth chorion CTBs remains unclear.

Several pieces of evidence suggest that the smooth chorion is not simply a vestigial structure. First, the intact CTB layer contains proliferating cells and is maintained until term (Yeh et al., 1989; Benirschke et al., 2006, 1979; Garrido-Gomez et al., 2017). Second, the histological heterogeneity among smooth chorion CTBs suggests functional distinctions. Yeh et al. (1989) characterize two distinct populations of vacuolated and eosinophilic CTBs. Vacuolated CTBs were positive for placental lactogen and placental alkaline phosphatase, while the eosinophilic subpopulations was not. Both populations were rich in keratin and neither had the known characteristics of villous CTBs. Bou-Resli et al. (1981) also note high levels of variation among CTBs in the smooth chorion and the existence of a vacuolated population. A more molecular characterization was carried out by Garrido-Gomez et al. (2017), which demonstrated heterogeneity of ITGA4 and HLA-G expression, markers previously associated with stemness and invasion, respectively (Genbacev et al., 2016, McMaster et al., 1995). This study also uncovered an expansion of the smooth chorion in cases of severe pre-eclampsia, along with a disease-specific gene expression pattern. In sum, these results suggest the smooth chorion CTBs are a heterogeneous and dynamic collection of cells with important functions in development and disease.

Single cell RNA-sequencing (scRNA-seq) has emerged as the standard for transcriptional characterization of complex organs. This methodology was previously applied to the placenta, but with a focus on the maternal-fetal interface, specifically the chorionic villi and basal plate (Liu et al., 2018; Suryawanshi et al. 2018; Vento-Tormo et al., 2018). Recently, scRNA-seq was used to profile the smooch chorion at term (Pique-Regi et al., 2019; Pique-Regi et al., 2020; Garcia-Flores et al., 2022). However, in Pique-Regi et al., 2019, only 132 CTBs were identified in the smooth chorion out of 29,921 cells (0.44%) collected from this region. A comparable number of CTBs in the SC were recovered in Garcia-Flores et al., 2022, potentially reflecting a difficulty in capturing these cells at term. To better understand the differences in the cell types and functions of the two sides of the developing placenta, we applied scRNA-seq to matching samples of cells isolated from the villous and smooth chorion regions of human samples from mid to late in the second trimester. We used scRNA-seq to compare the composition and developmental trajectories of CTBs in the VC and SC. The data were validated and extended with functional studies to gain initial insights into the basis of differential migration of trophoblasts in each region. These results identified a novel smooth chorion-specific CTB population important for establishment of a protective barrier and the suppression of trophoblast invasion. In addition, these data represent a resource of CTB types, proportions, and gene expression at mid-gestation against which age related and pathogenic alterations can be measured.

## Results

### The transcriptional landscape of the villous and smooth chorion at mid-gestation

To understand the cellular composition of the smooth chorion, we isolated and profiled cells from both the VC and the SC regions of four second trimester human placentas spanning gestational weeks 18 to 24 (GW18-24) using single cell RNA-sequencing (Figure 1a). We chose to analyze second trimester samples because the maturation of the smooth chorion is complete but the inflammation and apoptosis associated with membrane rupture and parturition is absent (Benirshcke et al., 2006; Yuan et al., 2006; Yuan et al., 2008; Yuan et al., 2009; Figure 1a). SC and VC cells were isolated from each human placental sample allowing within and across patient comparisons. VC samples included cells isolated from floating and anchoring villi and areas surrounding the cell column, while most of the decidua (including spiral arteries) were dissected away. SC samples included the chorion and underlying stroma (mesenchymal and endothelial cells), but not the amnion and little of the neighboring decidua, which were also removed during dissection. CTBs were further enriched over stromal and immune cells during cell preparation as previously described (Garrido-Gomez et al. 2017 and in methods). The transcriptomes of the resulting cells were captured using the 10x Genomics scRNA-seq platform.

**Figure 1.**
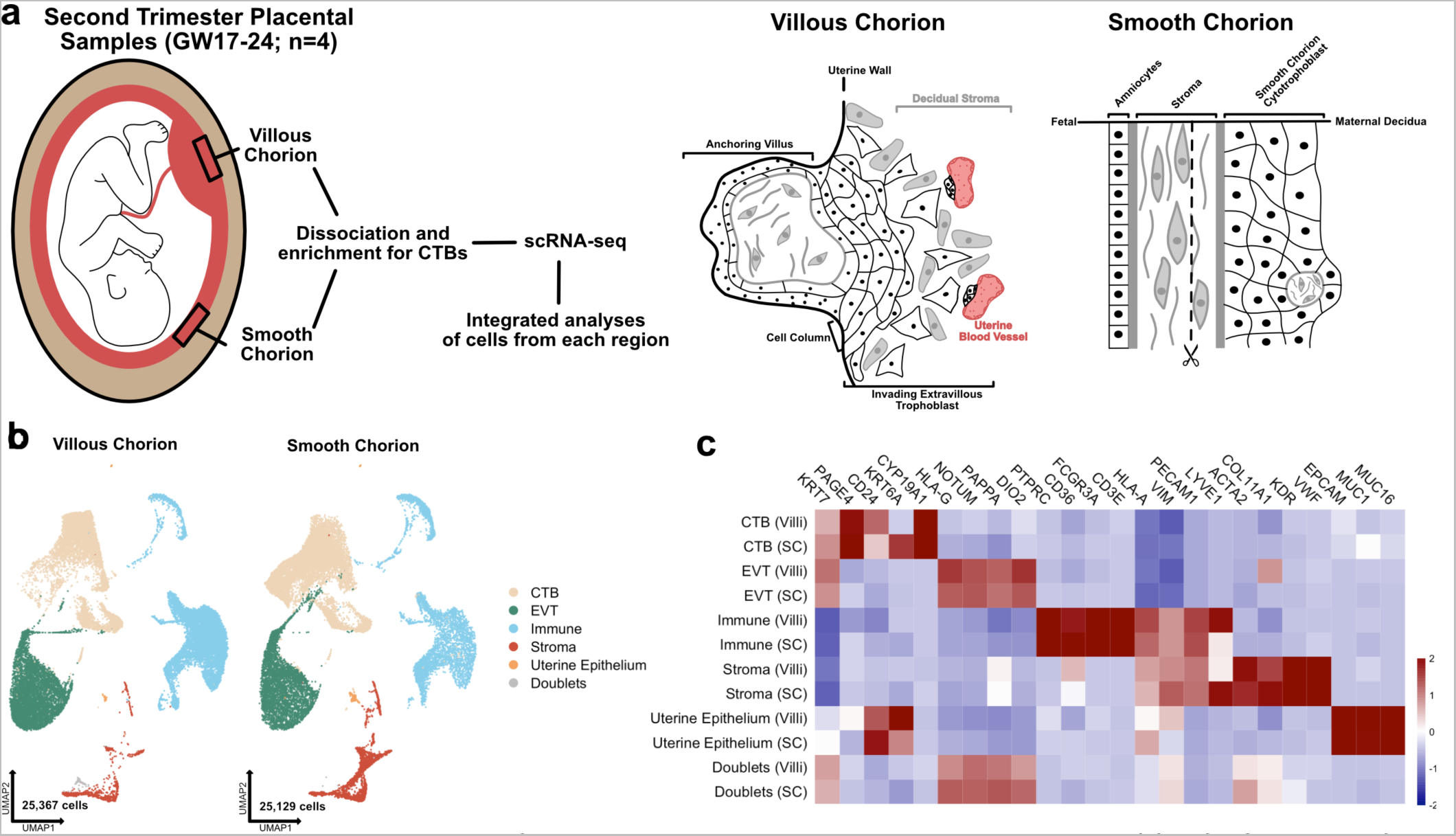
The transcriptional landscape of the villous and smooth chorion at mid-gestation. **(a)** Left - Schematic of the placenta at mid-gestation, highlighting the regions sampled, together with the methods used for cell isolation and characterization. Right – schematic of the cell types and their organization in each region. **(b)** UMAP projections of integrated samples, shown by region of origin (Left – VC, Right – SC), and colored according to broad cell type clusters. **(c)** Heatmap of the transcript expression of select cell identity markers across broad cell type clusters and regions. Values are scaled expression across the clusters of each region independently.

Each of the eight datasets (GW17.6, 18.2, 23.0, 24.0; VC and SC) were captured independently, then integrated computationally (Figure 1b; Stuart et al., 2019). We classified the cells of the integrated dataset into broad cell type clusters according to functional identities and annotated each by expression of canonical markers. CTBs were annotated as *KRT7*+, *HLA-G*^−^; EVTs as *KRT7*+, *HLA-G*+; immune cells as *CD45*+, *VIM*–; stromal cells as *VIM*+, *CD45*–; and the uterine epithelium by expression of *EPCAM*+, *MUC1*+ and *MUC16*+ (Figure 1c; Lee et al., 2016; McMaster et al., 1995; Vento-Tormo et al., 2018). Cells expressing exclusive markers of disparate cell types (co-expression of *KRT7, HLA-G, HLA-A, VIM, ACTA2*) were labelled as doublets and excluded from further analysis. The complete dataset used for further analysis contained 50,496 cells that passed quality control (between 500 and 6500 unique genes, fewer than 15% mitochondrial reads, doublets removed) [Figure 1-S1a; McGinnis et al., 2019]. Cells originating from each placental region (VC – 25,367 and SC – 25,129) and each placental sample (7,181-17,705 cells per sample) were well represented (Figure 1b; Figure 1-S1b and c). The number of cells in each broad cell type cluster demonstrated enrichment for CTBs, which represented more than 60% of the cells in the integrated dataset, as expected given the enrichment protocol (Figure 1-S1d). The sex of each fetus was inferred by assaying expression of *XIST* in trophoblast cells isolated from each sample (Figure 1-S2a), which allowed the assignment of cell types to either fetal or maternal origin (Figure 1-S2b).

Even though there was enrichment for CTBs, we still identified 14,805 immune cells and 3,883 stromal cells. We reclustered these immune and stromal cell subsets individually and compared them to previously published single cell analyses, allowing for the annotation of subtypes within each group (Figure 1-S3a-d; Figure 1-S4a-d; Figure 1-S5a-c; Figure 1-S7a-c)[Vento-Tormo et al., 2018, Pique-Regi et al., 2019]. Independent clustering analysis of VC and SC cells identified the similar populations as the integrated immune and stromal subsets and confirmed the relative proportions of cell identities across each region (Figure 1-S6a-d; Figure 1-S8a-d). All immune clusters robustly expressed *Xist* in each sample, identifying them to be of maternal origin (Figure 1-S2c). Almost two-fold more immune cells were recovered from the VC than the SC, although it possible that this change in proportion is an artifact of dissection and/or the CTB enrichment protocol (Figure 1-S5d and e). Comparing the immune cell types identified in each region revealed a higher proportion of macrophages in the VC as compared to the SC, which contained a greater proportion of NK/T cells (Figure 1-S5e; Figure 1-Source Data 1).

While few stromal cells were isolated in the preparations, sub-clustering still revealed a differential composition of fetal stromal cells between the VC and SC (Figure 1-S7a-c; Figure 1-Source Data 2). The majority of stromal cells recovered originated from the SC (2,941 compared to 942 from VC). These cells included lymphatic endothelium (Pique-Regi et al., 2019) and two largely SC-specific mesenchymal cell populations of fetal origin, Mesenchyme 1 and Mesenchyme 3 (Figure 1-S7c-e; Figure 1-S2b). These two clusters are marked by elevated expression of *EGFL6*, *DLK1*, and uniquely by expression of *COL11A1*, which is observed only in the SC (Figure 1-S7b and f). Interestingly, several canonical CTB support factors including HGF, WNT2, and RSPO3, were expressed in fetal stromal populations in both placental regions, suggesting shared requirements for WNT and MET signaling (Figure 1-S7g). Taken together these data demonstrate the identification of broad classes of CTBs, immune, and support cells from both the VC and SC regions of the human placenta.

### Identification of a smooth chorion-specific cytotrophoblast population

CTBs are the fetal cells that perform the specialized functions of the VC, and are required for normal fetal growth and development (Maltepe and Fisher, 2015; Turco and Moffett, 2019; Knöfler et al., 2019). To better understand the composition of CTBs in the SC versus the VC, we sub-clustered this population (*KRT7*+, *VIM-*, *CD45-*, *MUC1-*). The CTB subset is comprised of 29,668 cells with similar representation and cell quality control metrics across all eight samples (Figure 2-S1a, b, and c). This analysis identified 13 clusters including several CTB, EVT, and STB subtypes (Figure 2a; Figure 2-Source Data 1).

**Figure 2.**
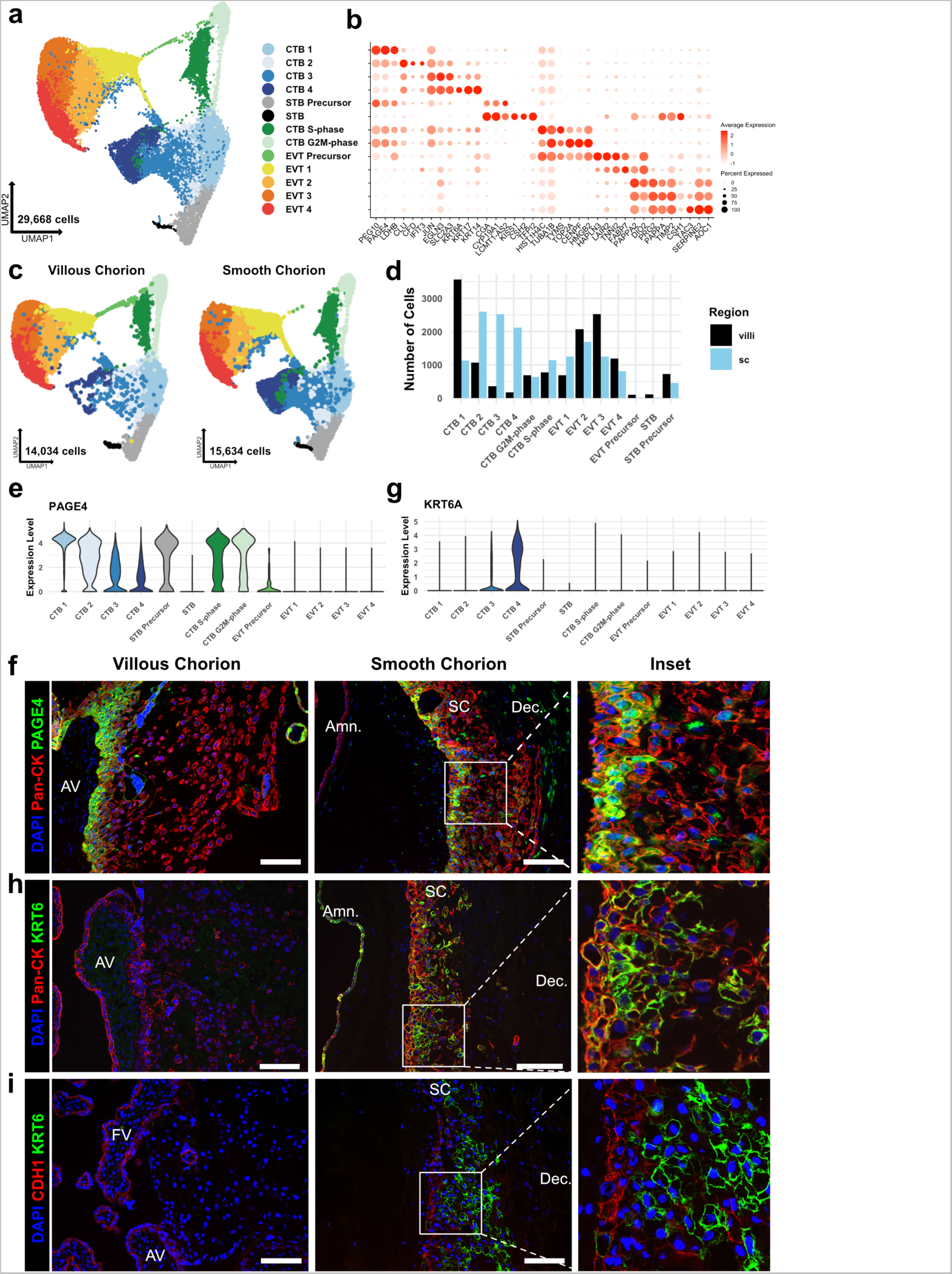
Identification of a smooth chorion-specific cytotrophoblast. **(a)** UMAP projection of subclustered trophoblasts (n=29,668). Colors correspond to the clusters at the right. **(b)** Dot plot showing average expression and percent of cells in each cluster as identified by the marker genes listed on the x-axis. The clusters are listed on the y-axis. **(c)** UMAP projection of subclustered trophoblasts from the VC (Left) or the SC (Right). Clusters and colors are the same as in panel a. **(d)** Quantification of the number of cells in each trophoblast cluster from each region. Cells from the VC are shown in black. Cells from the SC are shown in blue. **(e)** Violin plot of *PAGE4* transcript expression across all trophoblast clusters. (**f)** Immunofluorescence co-localization of PAGE4 with pan-cytokeratin (marker of all trophoblast) in the VC (Left) or SC (Right). **(g)** Violin plot of *KRT6A* transcript expression across all trophoblast clusters. **(h)** Immunofluorescence co-localization of KRT6 with pan-cytokeratin (marker of all trophoblast) in the VC (Left) or SC (Right). **(i)** Immunofluorescence co-localization of CDH1 and KRT6 in the VC (Left) or SC (Middle). High magnification inset is denoted by the white box (Right). For all images nuclei were visualized by DAPI stain; Scale bar = 100μm. Abbreviations: AV = Anchoring Villi; FV = Floating Villi; SC = Smooth Chorion epithelium; Amn. = Amnion; Dec. = Decidua.

Broad classes of trophoblast were annotated by established markers (STBs – *CGA*, *CYP19A1*, *CSH1*, *CSH2*; EVTs – *HLA-G, DIO2*; CTBs – *PAGE4*, *PEG10,* and no expression of EVT and STB markers) [Figure 2b and Figure 2-S2a; McMaster et al., 1995; Lee et al. 2016, Suryawanshi et al. 2018, Liu et al. 2018]. Comparison to previously published cell types in the VC and SC confirmed the identities of most clusters (Figure 2-S1d-g; Vento-Tormo et al., 2018; Pique-Regi et al., 2019). Two cell clusters showed high expression of canonical phasic transcripts, including *MKI67*, with the S-phase cluster denoted by expression of *PCNA* and the G2/M-phase cluster by expression of *TOP2A* (Figure 2-S2b; Tirosh et al. 2015). Both populations share gene expression with all clusters of CTBs, and therefore, were identified as actively cycling CTBs (Figure 2-S2a). No STB or EVT markers were identified in the cycling clusters as was expected due to the requirement for cell cycle exit upon terminal differentiation to these lineages (Lu et al. 2017, Genbacev et al., 1997).

CTBs separated into four clusters, CTB 1-4. CTB 1 cells highly expressed *PAGE4*, *PEG10*, and *CDH1* (Figure 2b, Figure 2-S2a, Figure 2-S4a), which have been shown to be canonical markers of villous CTB (Lee et al. 2016, Suryawanshi et al. 2018). Overall, CTB 2-4 were more transcriptionally similar to each other than to CTB 1, indicating a transcriptional program that was distinct from canonical villous CTBs (Figure 2-S2a and c). CTB 2-4 existed along a gradient of gene expression changes suggestive of various stages of a common differentiation pathway. However, each population expressed distinct transcripts corresponding to important proposed functions of the smooth chorion. CTB 2 cells highly expressed *CLU*, *CFD*, and *IFIT3*, suggesting roles in responding to bacterial or viral infection through the innate arm of the immune system (Thurman and Hollers., 2006; Liu et al., 2011). CTB 3 cells upregulated *EGLN3* and *SLC2A3*, indicating a HIF-mediated response to hypoxia and a switch toward glucose metabolism, likely as an adaptation to the decreased oxygen levels in the largely avascular smooth chorion region (del Peso et al. 2003, Maxwell et al., 1997). Finally, CTB 4 specifically expressed several cytokeratins — *KRT6A*, *KRT17*, and *KRT14*, found in many epithelial barrier tissues and important for maintenance of integrity in response to mechanical stressors (Karantza, 2011; Figure 2b, Figure 2-S2a). Previous analysis of the SC region at term using scRNA-seq identified fewer than 0.5% of trophoblasts as being CTB and did not identify a separate KRT6 expressing CTB population (Pique-Regi et al., 2019). Computational integration of the data from Pique-Regi et al., 2019 with the trophoblast subset confirmed CTB 3 and CTB 4 to be unique to this study (Figure 2-S1f and g). These results establish a previously underappreciated transcriptional diversity of CTB subpopulations.

Quantification of cells showed a strong regional bias in the number and proportion contributing to each CTB cluster (Figure 2c and d). In the VC samples, 53.8% of CTBs clustered in CTB 1 compared to 11.1% in the SC (Supplementary Table 1). In contrast, the SC had a much larger proportion of the CTB 2-4 clusters. In this region, 71.3% of CTBs were nearly equally distributed among CTB 2 (25.6%), CTB 3 (24.9%), and CTB 4 (20.8%). In the VC, only 24.1% of CTBs were found in the same clusters, with the majority in CTB 2 (16.1%). The contribution to cycling clusters was consistent across regions: 22.1% and 17.5% of CTBs in the VC and SC, respectively. The relative proportions of CTB 1-4 in the VC and SC were consistent across individual samples indicating that this difference was not driven by sample variability (Figure 2-S1b-c). Furthermore, independent clustering analysis of VC and SC cells identified the same populations as the integrated trophoblast subset. Importantly, independent clustering of each region did not identify CTB 4 in the VC or STB in the SC, suggesting CTB 4 and STB to be specific to the SC and VC, respectively. The small number of cells in the integrated dataset identified as VC CTB 4 (173 cells) or SC STB (14 cells) were likely an artifact of computational integration (Figure 2-S3a-f).

Next, we immunolocalized the protein products of genes that distinguished the subpopulations. At the mRNA level, *PAGE4* expression was highest in CTB 1 and decreased across CTB 2-4 (Figure 2b and e). In the VC, the CTB monolayer between the fetal stromal villous core and the overlying STB layer showed strong PAGE4 immunoreactivity, which diminished upon differentiation to EVT (Figure 2f - left). PAGE4 mRNA and protein expression matched that of known villous CTB marker CDH1 (Figure 2-S4a and b; Zhou et al., 1997). In the SC, the PAGE4 signal was strong in the epithelial layer directly adjacent to the fetal stroma and then decreased in cells distant from the basal layer, again matching CDH1 (Figure 2f - right, Figure 2-S4b - right). Both RNA expression and protein localization were consistent with CTB 1 cells existing on both sides of the placenta and occupying a similar niche.

Staining for KRT6, a marker highly enriched in CTB 4 cells (Figure 2g) showed a strikingly different result. Cells occupying the upper layers of SC epithelium showed a strong KRT6 signal, a pattern opposite to CDH1 (Figure 2h and i - right). KRT6 was absent from either the floating or anchoring villi of the VC (Figure 2h and i - left), although rare decidual resident KRT6 positive cells were identified in the VC region. (Figure 2-S4c, Figure 6-S1 - top). KRT6 isoforms, KRT6B and KRT6C, were not expressed, confirming KRT6A transcript and protein as highly specific markers of a CTB population found only in the smooth chorion (Figure 2-S4d). These data describe a novel subpopulation of CTBs unique to the SC, which going forward we term CTB 4 or SC-CTBs for smooth chorion-specific CTBs.

### A common CTB progenitor gives rise to STBs in the VC and SC-CTBs in the SC

Next, we investigated the developmental origin of the SC-CTBs. We performed RNA velocity analysis to predict the relationships between cells based on the proportion of exonic and intronic reads. These predictions are shown as vectors representing both the magnitude (predicted rate) and the direction of differentiation (Bergen et al. 2020). We first asked whether RNA velocity could recapitulate the well-established differentiation trajectories of trophoblasts in the VC (Knöfler et al., 2019; Turco and Moffett, 2019; Vento-Tormo et al. 2018). In accordance with previous results, RNA velocity projections identified CTB 1 as the root for three differentiation trajectories: self-renewal, differentiation to STBs, and differentiation to EVTs (Figure 3a). Cells at the boundary of the CTB 1 cluster showed differentiation vectors of high magnitude toward STB Precursors and upregulated canonical drivers of STB differentiation and fusion (*ERVW-1* and *ERVFRD-1*). These cells also expressed transcription factors (*GCM1* and *HOPX*) and hormones (*CSH1*) necessary for STB function (Figure 3-S1a; Baczyk et al., 2009; Mi et al., 2000; Blaise et al., 2003; Yabe et al., 2016).

**Figure 3.**
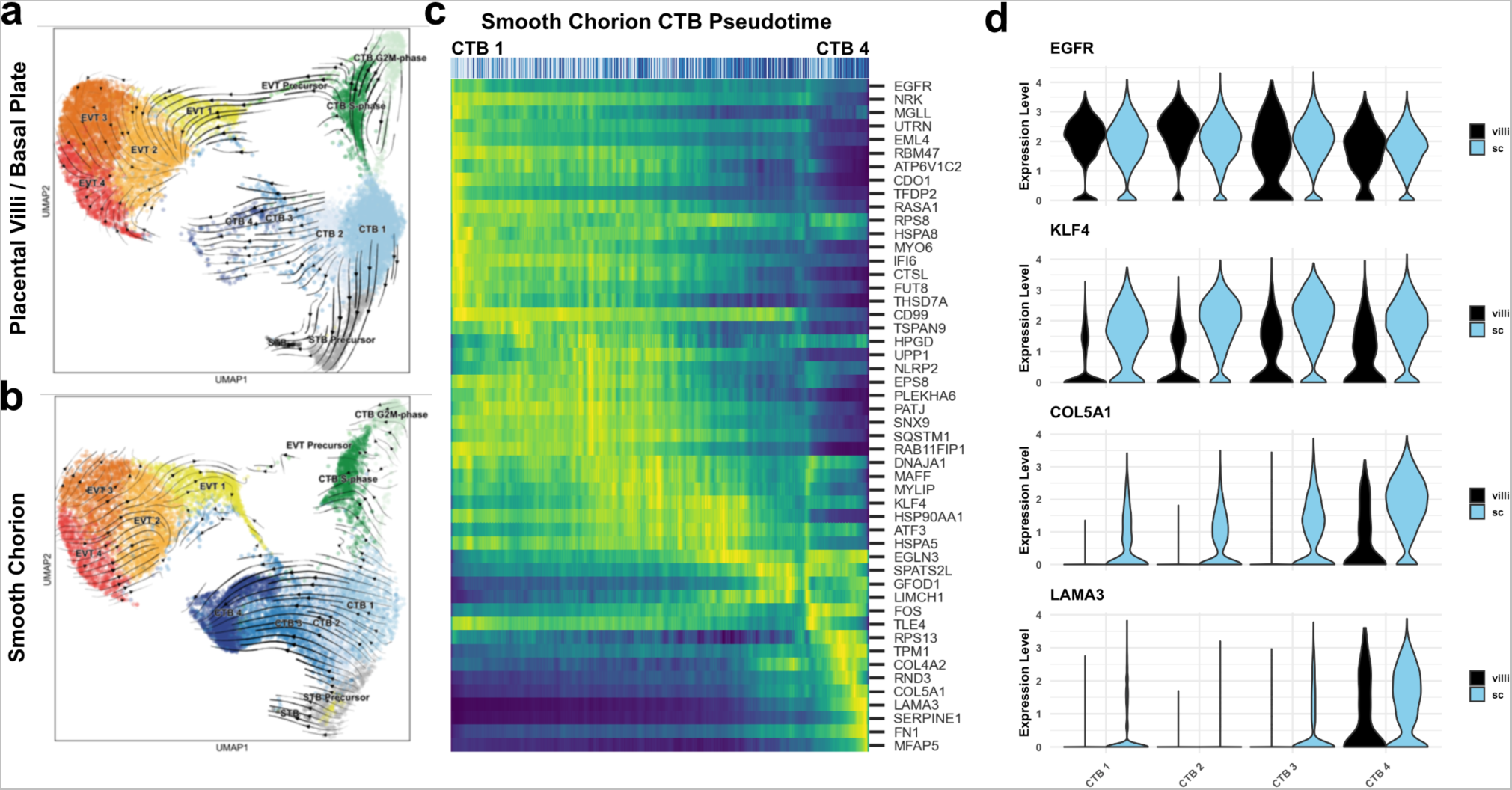
A common CTB progenitor gives rise to STBs in the VC and SC-CTBs in the SC. RNA velocity vector projections overlaid on to UMAP projections for trophoblast cells isolated from the **(a)** VC and **(b)** SC. Arrows denote direction and magnitude is represented by line thickness. **(c)** Pseudotime reconstruction of SC derived CTB 1-4 clusters from the scVelo dynamical model of latent time. Each column represents one cell. Cells at the left are clustered in CTB 1 and progress through CTB 2, 3, and 4 along the x-axis. Select genes that were the major drivers of the pseudotime alignment are shown on the y-axis. Expression ranged from dark blue (lowest) to yellow (highest). **(d)** Violin plots of select factors from (c) demonstrated shared or region-specific expression for genes associated with the CTB 4 differentiation trajectory.

In the SC, CTB 1 cells once again were identified as the root for differentiation. However, CTB 1 cells showed strong directionality and magnitude toward CTB 2-4 (Figure 3b). All cells in CTB 1-4 clusters displayed uniform directionality indicating a robust differentiation trajectory ending at CTB 4. High levels of transcriptional similarity between CTB 2-4 compared to CTB 1 suggested CTB 2 and 3 are intermediate states between CTB 1 and CTB 4 (Figure 2-S2c). Mitotic KRT6+ cells were identified, and while the interaction between the cell cycle and differentiation of SC-CTBs remains unclear, these data show that differentiation to SC-CTB does not require cell cycle exit, unlike STBs and EVTs (Figure 3-S1b). In contrast to the VC, we observed no velocity vectors with directionality toward the STB lineage from the CTB clusters in the SC samples. Further, the smaller number of STB precursors (460 cells) and STBs (14 cells) exhibited reduced expression of STB canonical markers such as ERVFRD-1 and GCM1, and notably, a near absence of ERVW-1 (Figure 3-S1a). These cells may be associated with ghost villi (Benirschke et al., 2006). In sum, these data show differential developmental trajectories for the CTB 1 cells in the SC and VC, with the former largely giving rise to SC-CTBs and the latter to STBs.

To identify the genes that were correlated with progression from CTB 1-4 in the SC, we used the velocity vector predictions to construct a pseudotemporal model of differentiation. All the cells in these clusters were plotted in one dimension from the least to the most differentiated according the pseudotime model (Figure 3c). Genes that were highly expressed at the start of the pseudotemporal differentiation included pan-trophoblast factors such as *EGFR*, which was expressed throughout all four CTB populations in both the VC and SC. Progression along the pseudotime trajectory identified regulators of cell fate and function, including the transcription factor *KLF4* and extracellular matrix (ECM) components *COL5A1* and *LAMA3* (Figure 3c-d); all demonstrated SC-specific expression. Elevated expression of ECM transcripts (*COL4A2*, *FN1*) and transcription factors responsive to cell contact and mechanical stress *(HES1*, *YAP1)* were coordinately upregulated, potentially highlighting the effects of the extracellular environment on fate specification (Figure 3-S1c). To identify differential signaling events that might regulate alternative paths of differentiation in the VC and SC, we used CellPhoneDB to predict receptor-ligand interactions between CTB clusters within each region (Figure 3-S2a-b; Efremova et al., 2020). This analysis identified BMP, Notch, and Ephrin signaling events specific to the SC region, which may help to determine cell fate and/or cell sorting within the SC trophoblast epithelium (Figure 3-S2b). Together, these data demonstrated that SC-CTBs originate from CTB 1 progenitors common to both the VC and SC. In the SC, instead of upregulating syncytialization factors such as *GCM1* and *ERVFRD-1*, CTB 1 progenitors upregulate transcription factors such as *KLF4*, *YAP1*, and *HES1*, which drive an epithelial cell fate in other contexts (Segre et al., 1999, Harvey et al., 2013; Rock et al., 2011).

### SC-CTBs express a distinct epidermal transcriptional program

Next, we sought a better understanding of the physiological functions of the SC trophoblast clusters. We performed gene ontology analysis as a summary of functional processes (Figure 4a, Figure 4-S1; Yu et al., 2012). We focused on the progenitor CTB 1 and terminally differentiated SC-CTBs as they showed enrichment for strikingly different functional categories. In CTB 1, we identified enrichment for WNT signaling, epithelial morphogenesis, and membrane transport, categories commonly associated with progenitors (Figure 4a, Figure 4-S1). We validated the activity of WNT signaling and the location of these cells by immunolocalization of non-phosphorylated CTNNB1. β-Catenin was localized to the most basal epithelial layer nearest to the stroma in both the VC and SC regions (Figure 4b), matching expression of the CTB 1 marker CDH1 (Figure 4-S2a). WNT signaling has an important role in the maintenance of villous CTBs *in vivo* and in the derivation and culture of self-renewing human trophoblast stem cells (Knöfler et al., 2019; Haider et al., 2018; Okae et al., 2018). We investigated proliferation of CTNNB1 expressing cells in both regions using KI67 as a mitotic marker. This revealed a similar percentage of KI67+ CTB 1 cells, suggesting similar proliferative capacity across regions (Figure 4b and c). We next analyzed regional differences within CTB 1. Gene ontology identified an enrichment for oxidative phosphorylation and epithelial signaling cues in VC CTB 1 cells (Figure 4-S2b). This is in direct contrast to SC CTB 1 cells that displayed elevated levels of hypoxia response genes (Figure 4-S2b-c). KLF4 was identified in the RNA velocity analysis as gaining expression from CTB1-4, but also showed greater expression in CTB1 in the SC compared to the VC (Figure 3d). In accordance with the mRNA expression data, KLF4 protein often co-localized to CTB1 in the SC, but was only observed in rare cells in the VC (Figure 4d). These data further support a similar location and function for CTB 1 in the SC and VC, with transcriptional and metabolic differences that presage distinct developmental trajectories.

**Figure 4.**
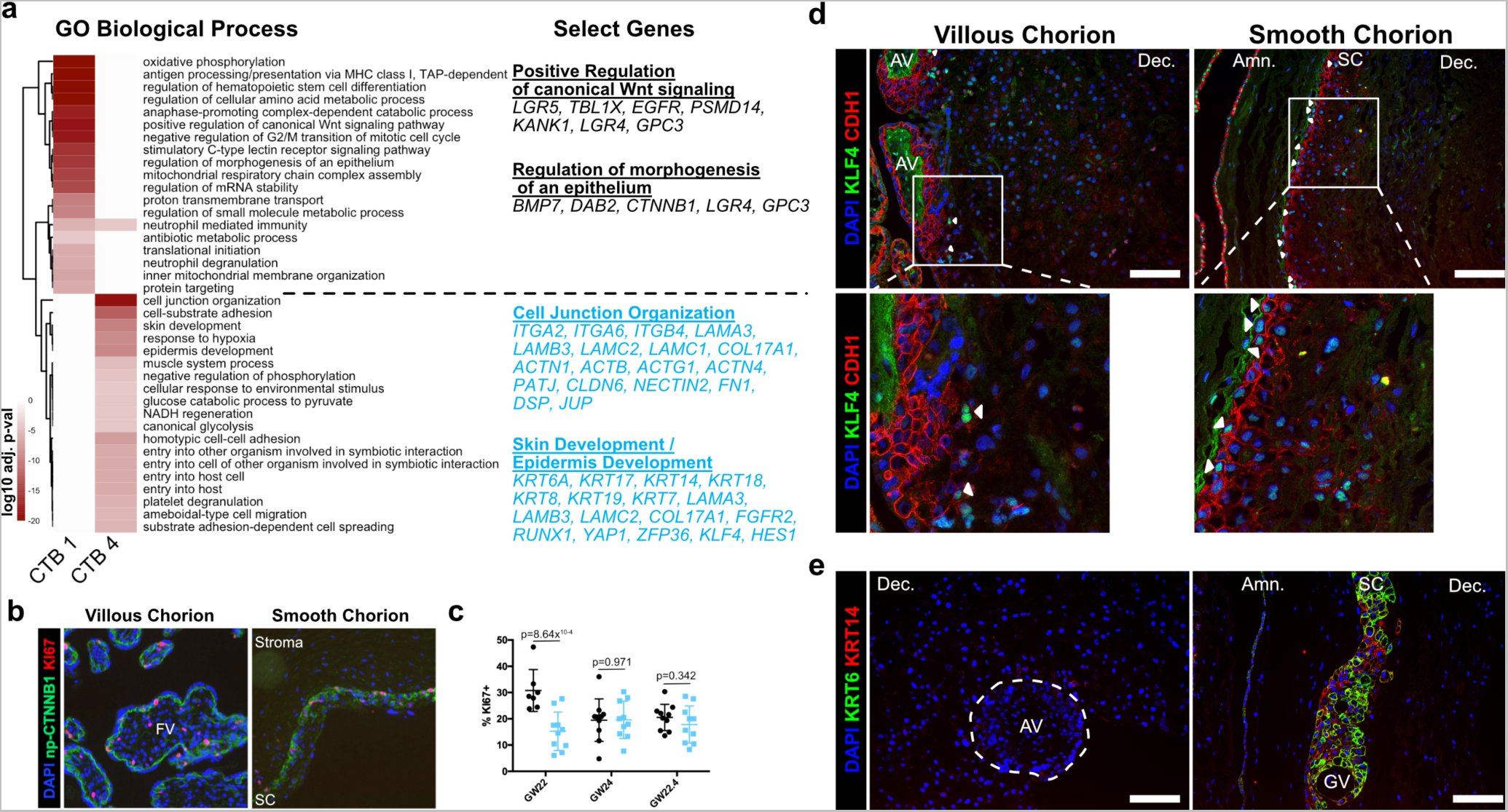
SC-CTBs express a distinct epidermal transcriptional program. **(a)** Heatmap of gene ontology analysis adjusted p-values. Dark red corresponds to the lowest p-values and white represents p-values greater than 0.0005. Ontology categories are organized by hierarchical clustering along the y-axis. Marker genes for each cluster were used as inputs for the analysis. A subset of genes in selected categories are listed at the right. Categories and genes enriched in CTB 1 or CTB 4 are in black or blue, respectively. **(b)** Representative immunofluorescence co-localization of non-phosphorylated CTNNB1 and KI67 in the VC (Left) and SC (Right). **(c)** Quantification of the percent of np-CTNNB1 cells with KI67 expression in each region in three placental samples. Each dot represents the percentage in one field of view (at least 7 per region per sample) as an estimate of mitotic cells per population. Percentages for the VC region are shown in black and the SC region in blue. **(d)** Immunofluorescence co-localization of CDH1 and KLF4 protein in the VC (Left) or SC (Right). Arrowheads denote CDH1+/KLF4+ cells. **(e)** Immunofluorescence co-localization of KRT14 and KRT6 protein in the VC (Left) or SC (Right). The outline of the Anchoring Villi (AV) is denoted by the white dashed line. For all images nuclei were visualized by DAPI stain; Scale bar = 100μm. Abbreviations: AV = Anchoring Villi; FV = Floating Villi; SC = Smooth Chorion epithelium; Amn. = Amnion; Dec. = Decidua; GV = Ghost Villi.

CTB 4 ontological analysis strongly supported important roles for these cells in the formation of a protective barrier. The greatest enrichment was for cell junction and cell substrate adhesion genes that included numerous integrin and laminin subunits as well as junctional proteins *PATJ*, *DSP*, and *JUP*. An enrichment for both skin and epidermal development correlated with upregulation of the ECM and junctional transcripts. These categories included many cytokeratins (*KRT6A, 7, 8, 14, 17, 18,* and *19*) and transcription factors (*KLF4*, *YAP1)* required for epidermal identity and maintenance (Segre et al., 1999; Schlegelmilch et al., 2011). The organization of the SC is reminiscent of stratified epidermal cells of the skin, with progenitors adherent to the basal lamina and more differentiated cells progeny forming the upper layers. In many tissues, the specific domains of cytokeratin expression correspond to stratified cell layers with different functions. We asked if this was also the case in the SC epithelium. Cytokeratins 7, 8, and 18 were expressed in all trophoblast regardless of region (Figure 4-S3a), but cytokeratin 6A, 14, and 17 displayed smooth chorion-specific expression that increased with differentiation towards CTB 4 (Figure 4-S3b). Immunofluorescence localization confirmed expression of KRT14 as specific to the CTBs in the SC and inclusive of all KRT6 expressing cells (Figure 4e). These data support a model of accumulated cytokeratin expression that begins with CTB 1 (*KRT7, KRT8, KRT18*), increases in CTB 2-3 (*KRT14, KRT17*), and culminates in CTB 4 (*KRT6A*). Together these data are consistent with a central role for the CTBs of the SC in establishing a protective epithelial barrier for the rapidly growing fetus.

Beyond forming a physical barrier, important placental functions include protection from bacterial and viral infections. We identified an enrichment for genes involved in antiviral response in SC-CTBs, and in SC cells more broadly. For example, IFITM3, a restriction factor preventing entry of viruses into cells, is highly expressed in CTBs from the SC compared to the VC (Figure 4-S4; Bailey et al., 2014; Spence et al., 2019). IFITM proteins have also been reported to inhibit syncytialization (Buchrieser et al. 2019), suggesting IFITM3 may also function to block differentiation of CTBs into STBs in the SC. Taken together, these data establish SC-CTBs as the building blocks and critical regulators of the SC barrier, responsible for both protection against physical forces and pathogen infection.

### EVTs of the VC and SC regions display distinct invasive activity but are transcriptionally similar

EVTs are the invasive trophoblasts of the placenta (Knöfler et al., 2019; Turco and Moffett, 2019; Red-Horse et al., 2004). While the EVTs of the VC migrate away from the villi, invade the decidua, and replace the endothelial lining of the uterine arteries, the EVTs of the SC adhere to the decidua and do not home to the maternal vasculature (Genbacev et al., 2015). To understand the basis for these differences, we compared EVT subpopulations isolated from the VC and SC. Expression of the canonical marker of EVTs, *HLA-G*, was similarly abundant among the cells isolated from both placental regions (Figure 5a and b). A greater number (VC - 6572; SC - 5021) and larger proportion (VC – 46.83%; SC – 32.12%) of CTBs from the VC expressed HLA-G as compared to the analogous population from the SC (Figure 5c). Consistent with the transcript expression data, immunolocalization of HLA-G showed strong staining of cells in both the VC and SC. However, VC derived HLA-G positive cells were found deep in the maternal decidua. In contrast, HLA-G positive cells in the SC remained in the epithelial layer (Figure 5b).

**Figure 5.**
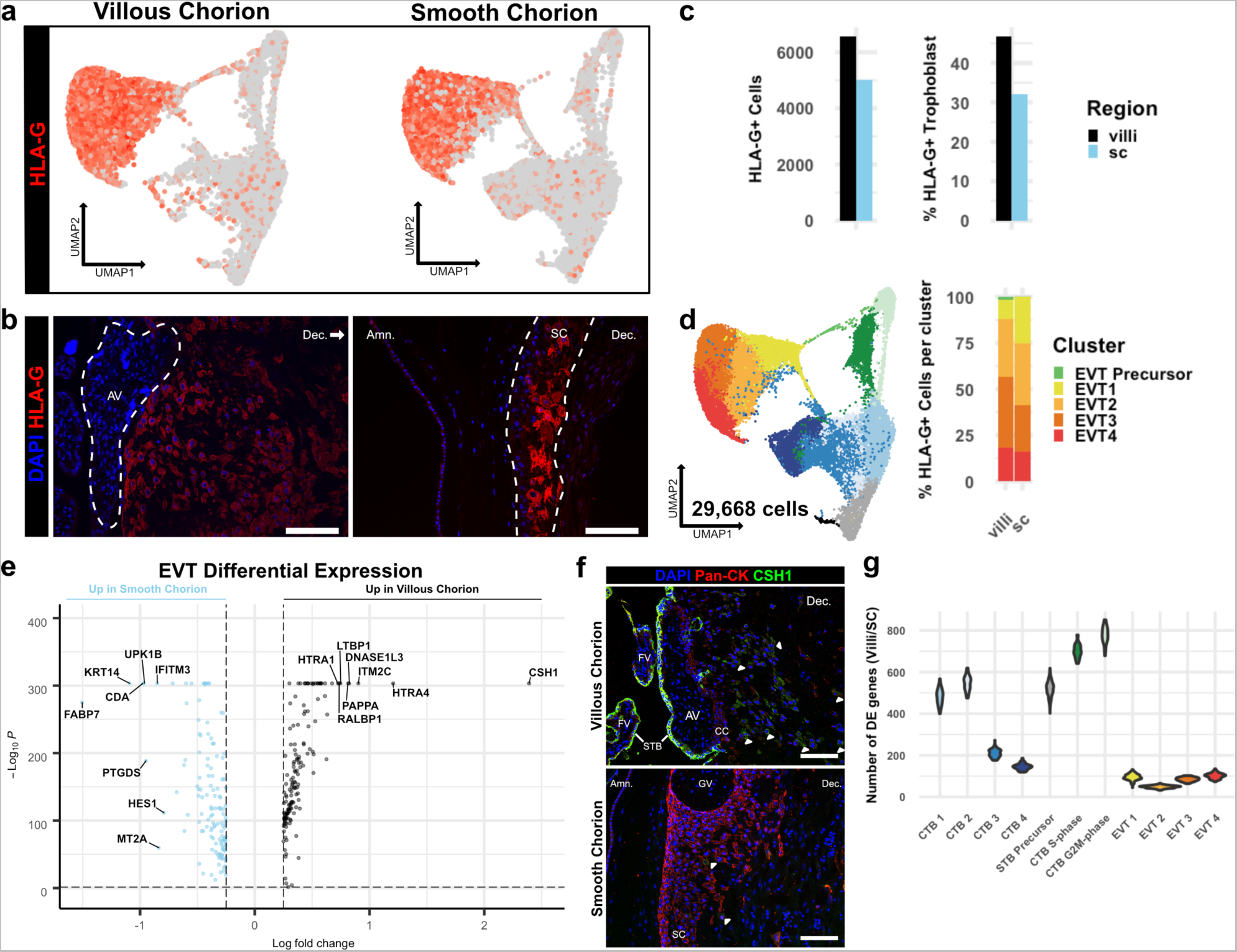
– EVTs of the VC and SC regions display distinct invasive activity but are transcriptionally similar. **(a)** Expression of *HLA-G* transcript per cell projected in UMAP space for the VC (Left) and SC (Right). Expression ranged from low in light gray to high in dark red. **(b)** Immunofluorescence localization of HLA-G in the VC (Left) and SC (Right). The Anchoring Villi (AV) are outlined in white (Left). The boundaries of the smooth chorion epithelium are denoted by the white lines (Right) **(c)** Quantification of the number of *HLA-G* expressing EVT (Left) and the percent of total trophoblast that express *HLA-G* (Right) for each placental region. **(d)** UMAP projection of all trophoblast cells including the EVT clusters (Left). The percent of EVT cells in each cluster from each region (Right). **(e)** Volcano plot of the differentially expressed genes between regions for all EVTs. All genes with a log fold change greater than an absolute value of 0.25 and a p-value of less than 0.05 were plotted. Those with greater expression in VC EVT are shown in black. Those with greater expression in SC EVT are shown in blue. **(f)** Immunofluorescence localization of CSH1 in the VC Top) and SC (Bottom). Arrowheads denote CSH1 expressing cells **(g)** Violin plots of the number of differentially expressed genes between 100 cells from each placental region within each cluster (100 permutations). Clusters with less than 100 cells per region were omitted due to the small sample size. For all images nuclei were visualized by DAPI stain; Scale bar = 100μm. Abbreviations: AV = Anchoring Villi; FV = Floating Villi; SC = Smooth Chorion epithelium; Amn. = Amnion; Dec. = Decidua; STB = Syncytiotrophoblast; CC = Cell Column.

**Figure 6.**
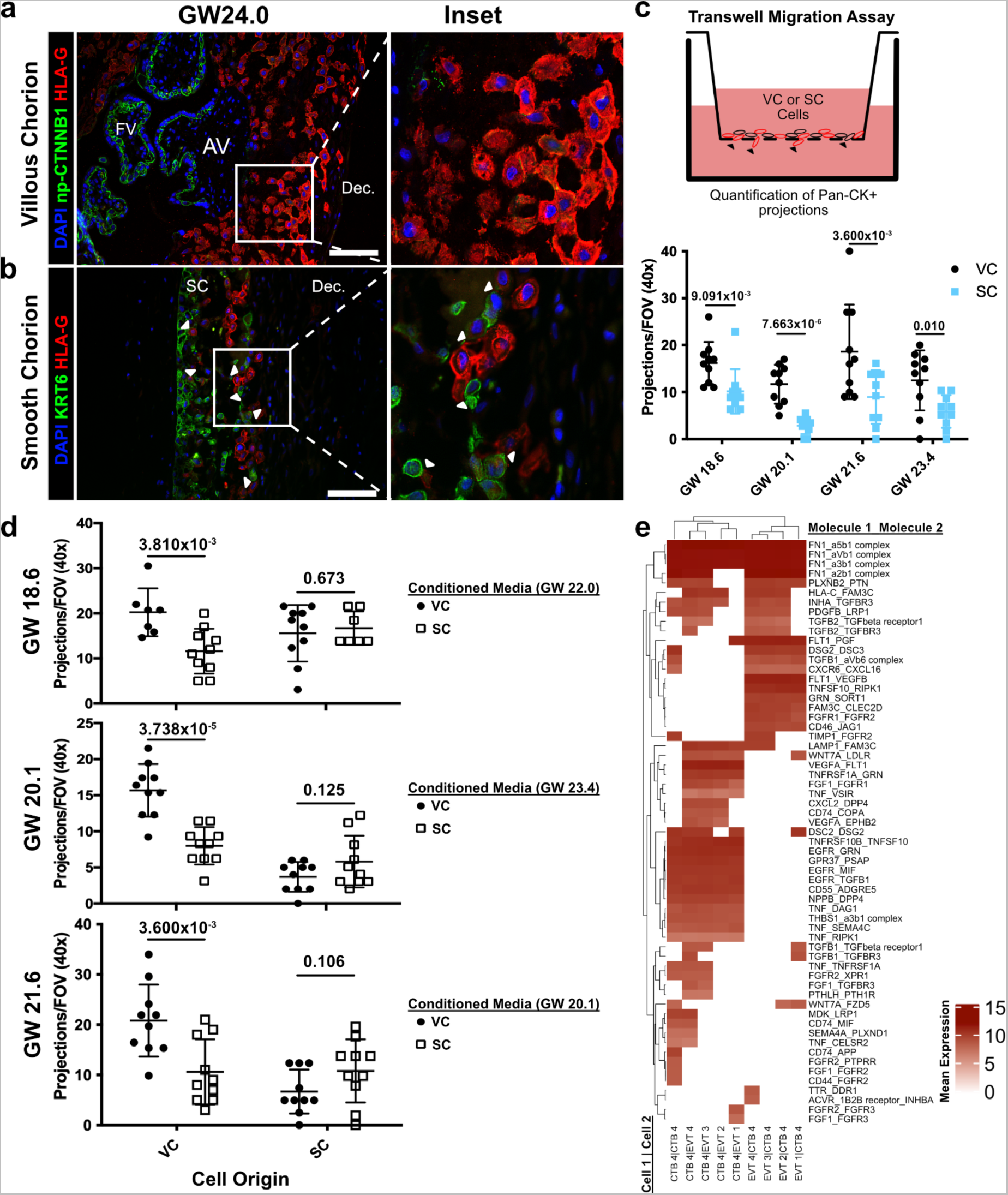
– CTBs of the SC inhibit EVT invasion. **(a)** Immunofluorescence co-localization of np-CTNNB1 and HLA-G in the VC. **(b)** Immunofluorescence co-localization of KRT6 and HLA-G in the SC. Arrowheads denote CTB and EVT interactions. **(c)** Schematic of the transwell invasion assay (top). Cells from either placental region were plated in the upper chamber of the transwell. After 39 hours of culture, the transwell membrane was fixed and stained with a Pan-cytokeratin antibody. The projections through the membrane are denoted by black arrowheads and quantified below. Results from the VC derived cells are shown in black, and the results from the SC derived cells are shown in blue. **(d)** Results from three biological replicates from each placental region cultured with conditioned medium from either VC or SC cells. The gestational ages of the plated cells is shown adjacent to the y-axis. The gestational ages of the cells from which conditioned medium was collected are noted in the legends at the right. The results for cells cultured with VC cell conditioned medium are denoted by black dots, and the results for those cultured in SC cell conditioned medium are denoted by open squares. P-values were determined by t-test and are listed above each comparison. **(e)** Predicted receptor-ligand interactions from CellPhoneDB. The strength of interaction is estimated by mean expression and is plotted in the heatmap. Receptor-ligand interactions and cell pairs are listed such that Molecule 1 is expressed by Cell 1 and Molecule 2 is expressed by Cell 2. For all images nuclei were visualized by DAPI staining ; Scale bar = 100μm. Abbreviations: AV = Anchoring Villi; FV = Floating Villi; SC = Smooth Chorion epithelium; Dec. = Decidua.

EVTs were further divided into clusters 1-4. No clusters were specific to the VC or SC, except for the small number of putative EVT Precursors (VC – 99 cells; SC – 8 cells) [Figure 5d, Figure 3a]. Previous work has classified EVTs on the villous side of the placenta as columnar, interstitial, or endovascular based on gene expression and location in the decidua (Tilburgs et al., 2015; Knöfler et al., 2019). The columnar subpopulation is believed to represent newly differentiated EVTs, which lie at the base of the columns that connect the anchoring villi to the uterine wall. Interstitial EVTs migrate through the decidua homing to maternal arteries, which they invade. The relationship between this subpopulation and endovascular EVTs that replace the maternal arterial endothelium is unclear, with evidence supporting endovascular EVTs arising either from interstitial EVTs and/or an independent origin (Harris et al. 2009; Red-Horse et al. 2004; Pijnenborg et al., 2011). We expected to capture both columnar and interstitial EVT populations, but few endovascular EVTs as the cellular preparations were largely devoid of arteries. We investigated the four clusters of EVT identified in our analysis in the context of the VC and the SC. Both placental regions contained cells from all clusters, although distinct regional biases were evident (Supplementary Table 1). The SC region contained almost twice as many EVT 1 cells as the VC (VC – 684; SC – 1256), whereas the VC contained almost twice as many EVT 2-4 as the SC (VC – 5789; SC – 3757). We next asked which of the EVT clusters corresponded to known EVT classifications, indicating maturation state or invasive capacity. Gene ontology analysis showed an enrichment for placental development and antigen presentation categories in EVT 1 (Figure 5-S1a). Representative marker genes for these categories in EVT 1 included several lineage-specific transcription factors including *GCM1*, *PPARG*, and *CEBPB* (Figure 5-S1b; Knöfler et al., 2019, Ferreira et al., 2016). In contrast, EVT 2-4 showed enrichment for extracellular structure and matrix organization, glycosylation, and peptidase activity. EVT clusters 3-4 specifically showed increased expression for transcripts of proteases such as *HTRA1*, *MMP2*, and *MMP11* (Figure 5-S1c). Based on the GO and specific gene enrichments, EVT 1 appeared most similar to columnar EVTs, while EVT 2-4 were consistent with interstitial EVTs that gain invasive capacity. The relative enrichment for EVT 1 in the SC (VC – 10.57% of EVT; SC – 25.05% of EVT) suggested an expansion in columnar-like EVTs at the expense of interstitial EVT. Conversely, the relative enrichment for EVT 2-4 in the VC region suggested an expansion of the invasive subpopulation (Figure 5d).

As almost 75% of EVTs from the SC region were in EVT clusters 2-4, we next asked if there were differences in gene expression between interstitial EVTs isolated from each region. This analysis identified relatively few differentially expressed genes, with the vast majority identified having a log fold change less than 1 (Figure 5-S2a; Figure 5-Source Data 1-4). The small number of differentially expressed genes with a fold change greater than 1 was typically identified across multiple EVT clusters, suggesting an overriding transcriptional program that might supersede subcluster designations. To identify genes associated with this program, we combined all EVT subclusters into one cluster for analysis (Figure 5e). Differential expression analysis revealed the hormone CSH1 and several proteases including HTRA1, HTRA4, and PAPPA to be expressed at significantly higher levels in the more invasive VC EVTs. However, of the most differentially expressed transcripts only CSH1, ITM2C, and DNASE1L3 were exclusive to the VC (Figure 5-S2b). Immunolocalization of CSH1 confirmed this result (Figure 5f). CSH1, also known as human prolactin, is a secreted hormone that signals through the receptor PRLR found on maternal cells. As such, the primary effect of CSH1 is likely non-cell autonomous, raising the possibility that the largest differences in EVTs between regions may not be in inherent invasive capacity, but rather their ability to communicate with the neighboring cells. Therefore, we asked whether differential EVT-stroma signaling interactions exist between the VC and the SC (Efremova et al., 2020). While this analysis underrepresents the contribution of the stroma due to its depletion by the trophoblast enrichment protocol, we identified enrichment for collagen-integrin interactions between EVT and stromal cells in the VC, which were not present in the SC (Figure 5-S3a). In contrast, SC specific EVT-stromal interactions included PDGFA and FN1 signaling (Figure 5-S3b). As we see few transcriptional changes in EVTs between regions, our data support a model by which differential signaling may create a stromal environment more amenable to EVT invasion.

We next asked whether the gene expression differences between the VC and SC were smaller for EVT clusters than for all other trophoblast clusters. Calculating the Spearman Correlation coefficient across each cluster, EVT 3, EVT 4, and CTB 1 (which we have established as common to both regions) were most similar and the only clusters with a coefficient greater than 0.75 (Figure 5-S2c). Since this analysis does not account for cluster heterogeneity or size, we quantified the number of differentially expressed genes between 100 randomly selected cells from each placental region within each cluster. Across 100 permutations, the number of differentially expressed genes between regions was the lowest for all four EVT clusters (Figure 5g). Together, these results showed that the EVTs of the VC and SC are surprisingly similar even though they have distinct levels of invasion *in vivo*.

### CTBs of the SC inhibit EVT invasion

Given the transcriptional similarity between EVT clusters originating from the VC and SC, we wondered if a non-cell autonomous program could explain their distinct migratory properties. Immunofluorescence co-localization of CTB 1, SC-CTBs, and EVTs showed striking differences in the relative positioning of the EVTs and CTBs in the two placental regions (Figure 6a and b). On the VC side, which lacks SC-CTBs cells, HLA-G+ EVTs were distant from the np-CTNNB1+ CTB 1 cell population as expected (Genbacev et al., 2015). By the second trimester they had migrated away from the cell column of the anchoring villi into the decidua (Figure 6a). In contrast, HLA-G+ EVTs in the SC were adjacent to and in physical contact with KRT6+ SC-CTBs (Figure 6b). The interactions between SC-CTBs and EVTs were numerous and widespread throughout the SC epithelium. Contacts between CTB 1 and EVTs in the SC were not observed and the VC contain only rare and irregular KRT6+ cells (Figure 6-S1). This close association suggested possible paracrine signaling events between SC-CTBs and EVTs, which might impact each cell type.

To test this theory, we attempted to recapitulate the differential invasive properties of VC and SC EVTs in a transwell migration assay. Briefly, these cells were enriched using the same protocol as described for the scRNA-seq experiments. The cells were plated on Matrigel coated transwell filters and cultured for 39 hours. Trophoblast projections that reached the underside of the filter, a proxy for invasion, were visualized by immunostaining with a pan-cytokeratin antibody (Figure 6c, Figure 6-S2). Consistent with the differences observed *in vivo*, there was greater invasion of VC as compared to SC trophoblast (Figure 6c). Next, we asked whether CTBs from the SC region secreted soluble factors that inhibited invasion. The invasion assays were repeated with VC cells cultured with conditioned medium from SC cells and vice versa. While VC conditioned medium had no impact on either VC or SC cells, SC conditioned medium significantly reduced migration of VC cells (Figure 6d). Neither conditioned medium impacted the density of VC or SC cells (Figure 6-S3). Therefore, a secreted factor from SC cells inhibited the migration of VC EVTs. Given that SC-CTBs were the only cell type unique to the SC side, it is highly likely that they produced the secreted factors that inhibit the invasion of EVTs to which they are juxtaposed. To identify which factors might be responsible for the repression of cellular invasion, we subset the predicted signaling interactions between CTB-EVT isolated from the SC (Figure 6-S4). We then subset this analysis for only those containing secreted factors, which identified numerous interactions between CTB 4 and EVTs including both canonical cell signaling pathways (TNFa, PGF, TGFB1, FGF1, and PDGFB) and modifiers of the ECM (FN1 and THBS1)[Figure 6e]. These data support a paracrine signaling mechanism by which CTB 4 cells restrict EVT invasion in the SC.

## Discussion

This study profiled the developing 2^nd^ trimester placenta, providing a high-resolution molecular description of the trophoblast in the SC and placing them in context of the trophoblast from the VC of the same placenta. In doing so, we addressed long standing observations concerning apparent similarities and differences between VC and SC trophoblast, deconstruct the trophoblast populations in the SC, and ascribe functions to the cells of this understudied region. Our study characterized a novel trophoblast population distinct from both extant CTB in the VC and from EVT found in either placental region. These trophoblasts reside only in stratified epithelium of the smooth chorion, express a specific cytokeratin (KRT6), and comprise at least 25% of all trophoblast in the region. Lineage reconstruction by RNA velocity suggested the progenitors for these smooth chorion-specific CTBs are a resident proliferative progenitor trophoblast with a transcriptional profile that matches villous CTB. We identified several potential regulators of the transition from a common villous CTB-like cell to SC-CTB, including epidermal transcription factors such as KLF4 and YAP1. These molecules along with region-specific signaling events drive an epidermal transcriptional program creating the building blocks for a barrier against both physical and pathogen stressors. Finally, we identified CTB-EVT interactions occurring only in the smooth chorion and provide evidence that paracrine signaling between these populations restricts trophoblast invasion.

Since the smooth chorion is created by a degenerative process, consensus has been that these cells are remnants of this process, making the smooth chorion a vestigial structure (Benirschke et al., 2006). However, several previous observations suggested this might be an oversimplification. First, these cells were noted to be proliferative, suggesting either cellular expansion or turnover (Benirschke et al., 2006). We confirmed the presence of proliferative trophoblasts in the smooth chorion. Each cycling CTB cluster expressed markers of CTB 1-4 in both regions (Figure 2-S2a) and both CTB 1 (Figure 4b-c) and SC-CTBs (Figure 3-S1b) also expressed KI67 protein. Proliferation of both the progenitor and differentiated cells suggested a need for expansion coordinated with the growth of the developing fetus. Second, the importance of the smooth chorion was suggested by the noted heterogeneity of the trophoblasts within this region (Yeh et al., 1989; Bou-Resli et al., 1981; Garrido-Gomez et al., 2017; Benirschke et al., 2006). In agreement, our scRNA-seq results showed the smooth chorion to be a complex tissue with several trophoblast types. We identified, by differential transcription and function, at least two distinct CTB cell types, columnar and interstitial EVT, and a small number of STB Precursors. As discussed, all CTB and EVT populations either matched those found in the VC or had new region-specific functions. The only evidence of degeneration was the STB Precursor population within the smooth chorion, which lacked expression of both *ERVW-1* and *ERVFRD-1*, the fusogens necessary to form STB (Mi et al., 2000; Blaise et al., 2003; Liu et al., 2018). In sum, these studies suggested that the SC is a complex and functional tissue, not simply the remnant of villous degeneration.

We focused on CTB 4, a novel population found exclusively in the SC, in the context of the unique functions of this placental region. CTB 4 were enriched for epidermal, skin development, and antiviral gene categories suggesting concerted roles in the creation of a barrier against external forces and pathogens. Previous studies have noted that CTBs residing in the SC have high levels of keratin expression and are associated with extensive ECM deposits rich in laminin, collagen, and fibronectin (Yeh et al., 1989; Bou-Resli et al., 1981; Benirschke et al., 2006). Among all smooth chorion CTB populations, but most notably in CTB 4, we identified the upregulation of ECM components, including *LAMA3*, *COL5A1* and *FN1*. Expression of ECM molecules in the SC is essential to protect against premature rupture of the fetal membranes, but its deposition has previously been ascribed to the stromal cells of the chorion (Parry and Strauss, 1998). Our results suggested that the trophoblasts of the SC create a specific ECM environment distinct from other placental regions. This altered composition likely has wide-ranging effects on trophoblast fate, gene expression, and activity. It will be interesting to explore the impact of advancing gestational age on the SC epithelium, especially with respect to the ECM. Presumably, cellular and structural changes precede the normal process of membrane rupture prior to delivery at term. Whether premature rupture of the membranes phenocopies these events or is a unique process is an important open question.

Pathogen defense is another important function of the placenta. Infections of the chorion and amnion (chorioamnionitis) often result in adverse outcomes for mother and fetus, primarily preterm birth (Romero et al., 2014). We identified specific expression of the antiviral gene IFITM3 in CTB 4. IFITM3 expression is particularly interesting due to the recent finding that it blocks STB fusion mediated by the endogenous retroviral elements, ERVW-1 and ERVFRD-1. Therefore, IFITM3 expression in SC-CTBs may have a dual role – restricting viral entry into the cell and blocking the formation of STBs in the SC. Its expression also provides a mechanism by which the lack of STBs in the smooth chorion is maintained after degradation of the villi is complete.

A distinct feature of the CTBs of the SC was remodeling of the cytokeratin network coordinated with differentiation from CTB 1 to SC-CTBs (Figure 2g, 1Figure 4-S3b). The progressive expression of KRT14, KRT17, and KRT6A is reminiscent of the cytokeratin code found in the epithelial cell layers of many tissues (Karantza et al., 2011). KRT14 is recognized as a marker of all stratified epithelium including subsets of basal progenitor and stem cells in several tissues, usually alongside KRT5 (Moll et al., 1982; Nelson et al., 1983; Rock et al., 2009). While we did not identify expression of KRT5 in any trophoblast, the KRT14 expressing proliferative cells of the SC fit the profile of stratified epithelia in other locations. The roles of KRT17 and KRT6 are less clear. Most knowledge about the function of these molecules comes from the epidermis in the context of injury and disease. KRT6 and KRT17 are upregulated rapidly upon epidermal injury and are expressed through the repair phase, each contributing specific functions. (Takahashi et al., 1998; McGowan and Coulombe, 1998). KRT17 promotes proliferation and increases in cell size through Akt/mTOR and STAT3, respectively (Kim et al., 2006; Yang et al., 2018). KRT6 is a negative regulator of cell migration through the inhibition of Src kinase and associations with myosinIIA (Wong and Coulombe, 2003; Rotty and Coulombe, 2012; Wang et al., 2018). Finally, this molecule promotes expression of Desmoplakin and the maintenance of desmosomes, the latter, a long-established feature of SC trophoblasts (Bou-Resli et al., 1981; Bartels and Wang, 1983; Benirschke et al., 2006). Taken together, these observations are consistent with data on SC-CTBs. We found no evidence of their invasion, but did identify frequent interactions with the ECM, CTBs, and EVTs. The coordinated expression of these cytokeratins and the transcriptional profile writ large, suggested the SC has the properties of a highly specialized epidermis with a robust proliferative capacity and strong cohesive properties, but lacking migratory or invasive behavior.

As previous work has demonstrated or assumed, the majority of trophoblast identified are represented in both the VC and SC (Benirschke et al., 2006; Garrido-Gomez et al., 2017; Pique-Regi et al., 2019). We identified two cell types with striking transcriptional similarity between regions, CTB 1 and EVT. CTB 1 cells in the VC matched canonical villous CTB in transcriptional profile, activity, and niche. Surprisingly, we identified a similar population in the SC. They are localized to an epithelial sheet juxtaposed to the fetal stroma, separated by a thin basal lamina. These cells are supported by similar signaling pathways in both regions, including HGF and WNT (Figure 1-S7g; Dokras et al., 2001; Okae et al., 2018; Zhou et al., 2002). The similarities in location and growth factor requirements demonstrate that a niche similar to the villous trophoblast membrane, exists in the SC. Additionally, we identified CTB 1 as the progenitor population for SC-CTB, and possibly for EVT in the SC. Thus, CTB 1 in both regions function as multipotent progenitors. Despite the many similarities, the developmental trajectories emerging from CTB 1 differ in a region-specific manner. At present, it is unclear whether CTB 1 cells in both regions originally have the same potential or if fate restriction occurs as the fetal membranes form. Future experiments will need to functionally address whether CTB 1 from the VC can be coerced to generate SC-CTBs and if CTB 1 from the SC can efficiently generate STBs and invasive EVTs. Multipotent trophoblasts have been generated from first trimester smooth chorion cells, however, these cells have a distinct transcriptional profile from the second trimester cells we characterized (Genbacev et al., 2016).

Similarities between the EVT of each region, as defined by contact with the decidua, has been documented in observational, molecular, and most recently, transcriptomic studies (Benirschke et al., 2006, Pique-Regi et al., 2019). Our data confirmed a strong correlation between gene expression for all EVT populations, regardless of regional identity. Based on previously established EVT subtypes, we identified a reduction in the proportion of interstitial EVTs and a concomitant increase in columnar-like EVTs of the SC. However, we did not find evidence of specific EVT subpopulations or intrinsic gene expression programs that would explain the differences in the depth of invasion of EVT in each placental region. However, we did uncover CTB-EVT interactions specific to the SC epithelium. Rather than physically separating as they do in the VC, CTBs and EVTs co-occupy the stratified epithelium of the SC, providing a contained environment for paracrine signaling, cell contact mediated signaling, and cell-ECM interactions that may impact trophoblast invasion. In this study we verified that soluble signaling factors from SC cells restrict the invasion of their counterparts from the VC. We can speculate on potential candidates that may function across multiple biological systems to restrict invasion. CTB 4 cells express high levels of TIMP1, TIMP3, and SERPINE1 (PAI-1), which inhibit the function and/or activation of MMPs necessary for trophoblast invasion (Fisher et al., 1989; Zhu et al., 2012). Addition of recombinant TIMPs or a function blocking antibody against plasminogen (whose processing is inhibited by SERPINE1) reduced trophoblast invasion *in vitro* (Lala and Graham, 1990). While these molecules are expressed by decidual cells, we identified their expression within the SC epithelium directly adjacent to EVTs. Another candidate regulator of invasion expressed by CTB 4 is TNFα. Treatment of trophoblasts with TNFα increases EVT apoptosis, decreases protease expression, increases SERPINE1 expression, and decreases invasion *in vitro* (Otun et al., 2011; Xu et al., 2011). These data suggest that SC-CTBs secrete multiple molecules that could decrease the survival and invasion of EVTs in the SC. In this study we explored the role of secreted factors, but SC-CTBs may also influence EVTs through cell contact mediated signaling and the deposition of an ECM distinct from the basal plate. Future experiments will be necessary to explore the contributions of these distinct signaling pathways and matrices.

In summary, this study provides a high-resolution molecular accounting of the trophoblast that occupy smooth chorion. By comparison with trophoblasts isolated from the VC of the same placentas, we identified key similarities and differences to better understand the molecular determinants of trophoblast function and activity in the SC. We characterized a novel CTB population, marked by KRT6A expression, that is central to three key functions of the SC: formation of an epidermal-like barrier, blockage of aberrant STB differentiation, and restriction of EVT invasion. These data provide a better understanding of molecular and cellular pathways that control human placental development and against which disease related changes can be identified and therapeutic targets discovered.

## Key Resources Table

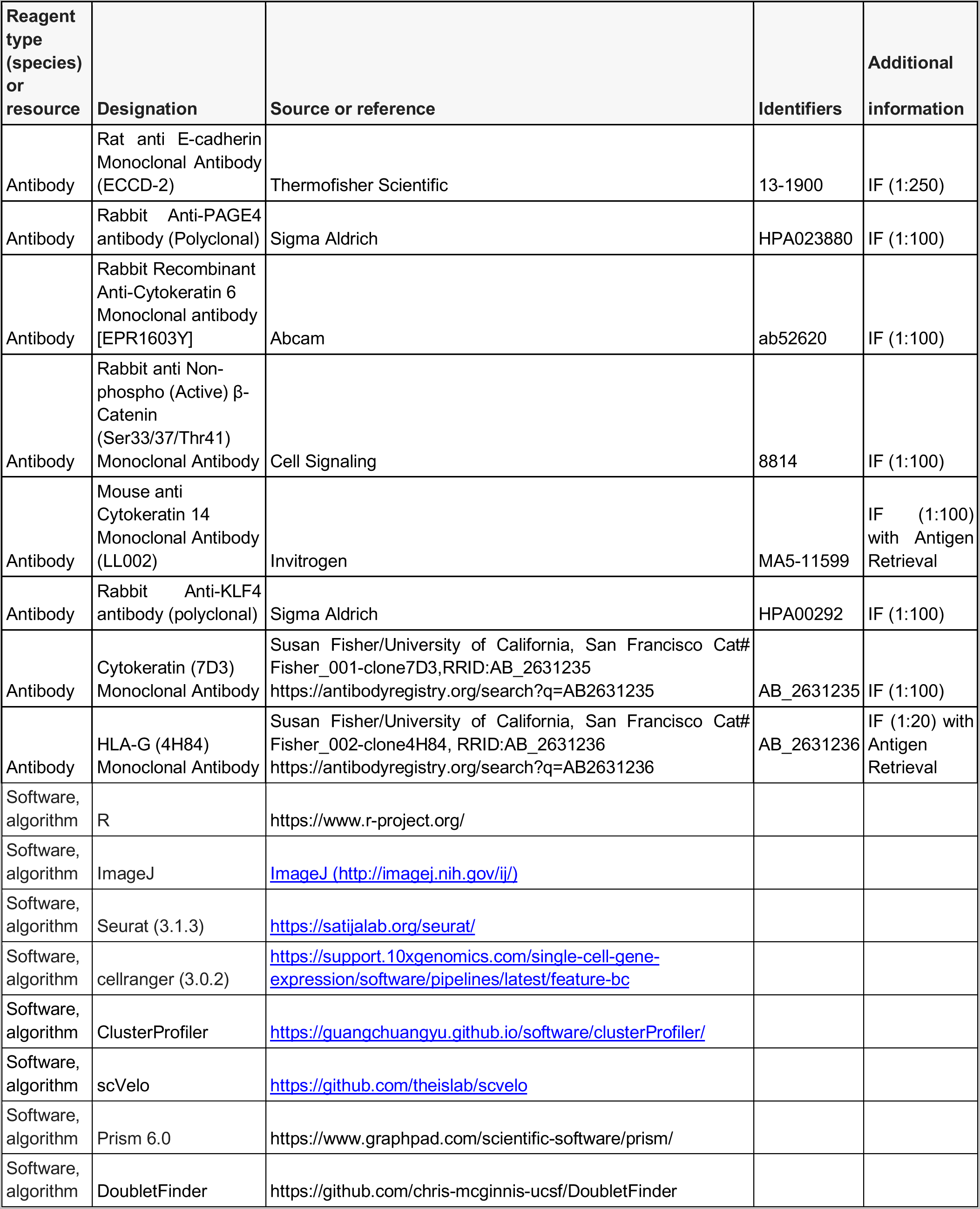

### Methods

#### Tissue Collection

The University of California, San Francisco (UCSF) Institutional Review Board approved this study. All donors gave informed consent. All samples are from elective terminations between gestational age of 17 weeks and six days and 24 weeks and zero days.

#### Cellular Isolation of VC trophoblast

Trophoblast were isolated from both floating and anchoring villi dissected from 2^nd^ trimester human placentas. Trophoblast were isolated according to previously published protocols (Fisher et al., 1989; Kliman and Feinberg, 1990). Briefly, cuts were made at the base of each cotyledon near the chorionic plate and the entire villous tree up to and including the basal plate was taken for study. The decidua was removed from the basal plate side and the remaining villous tree was dissected and dissociated. The resulting floating and anchoring chorionic villi were washed in cold phosphate-buffered saline (PBS Ca2+ and Mg2+ free), dissected into 2-4 mm pieces, and filtered through a 1 mm mesh strainer to remove small pieces of tissue. CTBs were isolated from the tissue pieces by first removing the outer syncytiotrophoblast layer by collagenase digestion (Sigma-Aldrich; C-2674). Next, CTBs were dissociated by sequential enzymatic digestion [trypsin (Sigma-Aldrich; T8003; twice) and collagenase]. Finally, CTBs were purified by Percoll density gradient centrifugation. Single cells were visually inspected for quality, counted using a hemacytometer, and immediately collected for single cell RNA-sequencing or for culture experiments.

#### Cellular Isolation of SC trophoblast

Trophoblast were isolated according to previously published protocols (Garrido-Gomez et al., 2017). Briefly, the fetal membranes were washed with phosphate-buffered saline (PBS Ca2+ and Mg2+ free) supplemented with 1% penicillin-streptomycin (10,000 units/ml penicillin; 10,000 μg/ml streptomycin), 0.003% fungizone (stock solution of 250 mg/ml) and 1% gentamicin. Next, the amnion and smooth chorion were manually separated and the amnion discarded. Next, the decidua parietalis was removed and discarded. The smooth chorion CTB layer was then minced into small pieces (2-4 mm) and dissociated by sequential enzymatic digestion. First, the tissue pieces were incubated in PBS (10 ml/g of tissue) containing 3.5 mg collagenase, 1.2 mg DNase, 6.9 mg hyaluronidase and 10 mg bovine serum albumin for 15-30 min. The supernatant was then discarded. Next, the tissue was incubated for 20-40 min in PBS containing trypsin (6.9 mg trypsin, 20 mg EDTA, 12 mg DNase per 100 ml; tissue weight: dissociation buffer volume=1:8). The enzyme activity was quenched by adding an equal volume of media containing 10% FBS. The cell suspension was filtered through a 70 μm sterile strainer and centrifuged at 1200 g for 7 min. A second collagenase digestion was performed by adding a 7x volume of the collagenase digestion buffer (see above), calculated on the basis of the weight of the cell pellet, followed by another incubation for 15-30 min. The cell suspension was then collected again by centrifugation. The cell pellets from the trypsin and second collagenase digestions were combined and purified over a Percoll density gradient centrifugation. Single cells were visually inspected for quality, counted using a hemacytometer, and immediately collected for single cell RNA-sequencing or for culture experiments.

#### Single Cell RNA-Sequencing and Analysis

To capture the transcriptome of individual cells we used the Chromium Single Cell 3’ Reagent V3 Kit from 10X Genomics. For all samples 17,500 cells were loaded into one well of a Chip B kit for GEM generation. Library preparation including, reverse transcription, barcoding, cDNA amplification, and purification was performed according to Chromium 10x V3 protocols. Each sample was sequenced on a NovaSeq 6000 S4 to a depth of approximately 20,000-30,000 reads per cell. The gene expression matrices for each dataset was generated using the CellRanger software (v3.0.2 - 10x Genomics). All reads were aligned to GRCh38 using STAR. (https://support.10xgenomics.com/single-cell-gene-expression/software/pipelines/latest/advanced/references). The counts matrix was thresholded and analyzed in the package Seurat (v3.1.3). Cells with fewer than 500 or greater than 6000 unique genes, as well as all cells with greater than 15 percent mitochondrial counts, were excluded from all subsequent analyses. Doublet detection was performed for each sample using DoubletFinder and all doublets were excluded from analysis. For each sample, counts were scaled and normalized using ScaleData and NormalizeData, respectively, with default settings FindVariableFeatures used to identify the 2000 most variable genes as input for all future analyses. PCA was performed using RunPCA and significant PCs assessed using ElbowPlot and DimHeatmap. Dimensionality reduction and visualization using UMAP was performed by RunUMAP.

Integration of each timepoint into one dataset was performed using FindIntegrationAnchors and IntegrateData, both using 20 dimensions (after filtering each dataset for number of genes, mitochondrial counts, and normalizing as described above). Data scaling, PCA, selection of PCs, clustering and visualization proceeded as described above using 30 PCs and a resolution of 0.6.

To generate the trophoblast, stroma, and immune cell subsets, the respective clusters were subset from the integrated dataset using the function SubsetData based upon annotations from marker genes identified by FindAllMarkers. After subsetting, counts were scaled and normalized using ScaleData and NormalizeData, respectively, with default settings FindVariableFeatures used to identify the 2000 most variable genes. Differentially expressed genes for each integrated dataset were identified using FindAllMarkers.

#### scVelo

RNA velocity analysis was applied to the entire conglomerate dataset using Velocyto to generate spliced and unspliced reads for all cells. This dataset was then subset for the trophoblast dataset introduced in Figure 2. The scVelo dynamical model was run with default settings and subset by each timepoint.

#### Differential expression between regions

The number of trophoblast cells in each cluster were downsampled to 100 cells from each region of origin and performed differential expression using FindMarkers function and repeated this for 100 permutations. The number of genes found to be significantly differentially expressed (adj. P-value <0.05) from each permutation are plotted. Clusters having less that 100 cells from each region were excluded from this analysis.

#### Gene Ontology Analysis

Gene Ontology analysis was performed with ClusterProfiler enrichGO function. The simplify function within this package was used to consolidate hierarchically related terms using a cutoff of 0.5. Terms were considered significantly enriched with an adjusted P-value of less than 0.05.

#### Immunofluorescence Staining

Placental tissue for cryosectioning was fixed in 3% PFA at 4**°**C for 8hrs, washed in 1x PBS, then submerged in 30% sucrose overnight at 4**°**C prior to embedding in OCT medium. Placental tissue in OCT was sectioned at 5μm for all conditions. In brief, slides were washed in 1x PBST (1x PBS, 0.05% Tween-20), blocked for 1hr (1x PBS + 5% donkey or goat serum + 0.3% TritonX), incubated in primary antibody diluted for 3 hours at room temperature (or overnight at 4**°**C), washed in 1x PBST, incubated in secondary antibody (Alexafluor 488 or 594) for 1 hour at room temperature, incubated in DAPI for 10min at room temp, washed in 1x PBST, and mounted and sealed for imaging. Any antigen retrieval was performed prior to the blocking step by heating the slides in a 1x citrate buffer with 0.05% Tween-20 at 95C for 30 minutes. All antibodies and the dilutions are listed in the key resources table. All immunofluorescence staining was performed in n=3 biological replicates (3 distinct placentas) and representative image is shown.

#### Transwell Invasion Assay

24-well plate transwell inserts with an 8 μM polycarbonate membrane (Corning Costar 3422) were coated with 10μL of Matrigel (growth factor-containing, Corning Corp, USA) diluted 1:1 in Serum Free Media (95% DME H-21 + Glutamine, 2% Nutridoma (mostly β-D xylopyranose), 1% Pen Strep, 1% Hepes, 0.1% Gentamycin). Cells from each region of the placenta were isolated as described above and then plated in the upper well of the transwell insert at a density of 250,000 cells in 250μL of Serum Free Media, with 1mL of Serum Free Media (or conditioned media) in the well below the insert. The cells were then cultured for 39 hours. The cells in the transwell were then fixed in 3% PFA for 10min at 4C, permeabilized in ice cold methanol for 10min at 4C, then washed in 1x PBS, incubated in Pan-CK primary antibody for 3 hours at 37C, washed in 1x PBS, incubated with secondary antibody (Alexafluor 594) and DAPI for 1 hour at 37C, washed in 1x PBS, then mounted and sealed for imaging. All experiments were performed in n=3 biological replicates (3 distinct placentas). For conditioned media experiments n=3 biological replicates were analyzed (cells and conditioned medium derived from 3 distinct placentas) .

#### Transwell Invasion Assay Quantification

Transwell membranes mounted on slides were imaged at 40x magnification and the number of Pan-CK positive projections through the membrane counted. Normalization for changes in cell density across fields of view and culture conditions was performed by quantifying the DAPI positive area of each image and the number of projections normalized to the median across comparisons. Between 7-10 fields of view were quantified for each transwell membrane. P-values were determined by t-test.

#### Culture of placental cells and generation of conditioned media

Each well of a 24-well plate was coated with 20μL of Matrigel undiluted (growth factor-containing, Corning Corp, USA) prior to cell seeding. Cells from each region of the placenta were isolated as described above and then plated into wells of a 24-well plate at a density of 1x10^6^ cells in 1mL of Serum Free Media (95% DME H-21 + Glutamine, 2% Nutridoma (mostly β-D xylopyranose), 1% Pen Strep, 1% Hepes, 0.1% Gentamycin). The cells were cultured for 39 hours. After 39 hours in culture the media was removed and centrifuged at 2000g for 10mins to remove cellular debris. The supernatant was then removed, snap frozen in LN2, and stored at -80C. This media was then thawed and used as conditioned media.

## Data Availability

All sequencing data is available at the NCBI Gene Expression Omnibus GSE198373. Processed data are available as R objects at https://figshare.com/projects/Regionally_distinct_trophoblast_regulate_barrier_function_and_invasion_in_the_human_placenta/135191. Code to process all raw data and generate the datasets analyzed are available at https://github.com/marshbp/Regionally-distinct-trophoblast-regulate-barrier-function-and-invasion-in-the-human-placenta. Previously published datasets from Vento-Tormo et al., 2018, and Pique-Regi et al., 2019 are publicly available.

## Acknowledgements

We thank the members of the University of California – San Francisco National Center of Translational Research in Reproduction and Infertility for helpful comments during the design, execution, and publication of this project. We would also like to thank all members of the Blelloch and Fisher Labs for their comments and support. We specifically thank Ali San and Nasim Zeighami for their research contributions. We would also like to acknowledge our funding sources, the NIH Eunice Kennedy Shriver National Institute for Child Health and Human Development P50 HD055764 and NIH R37 HD076253.

## Supplementary Tables

Supplementary Table 1 - Number of cells captured from each region per cluster for the trophoblast dataset.

## Supplementary Figures

**Figure 1-S1.**
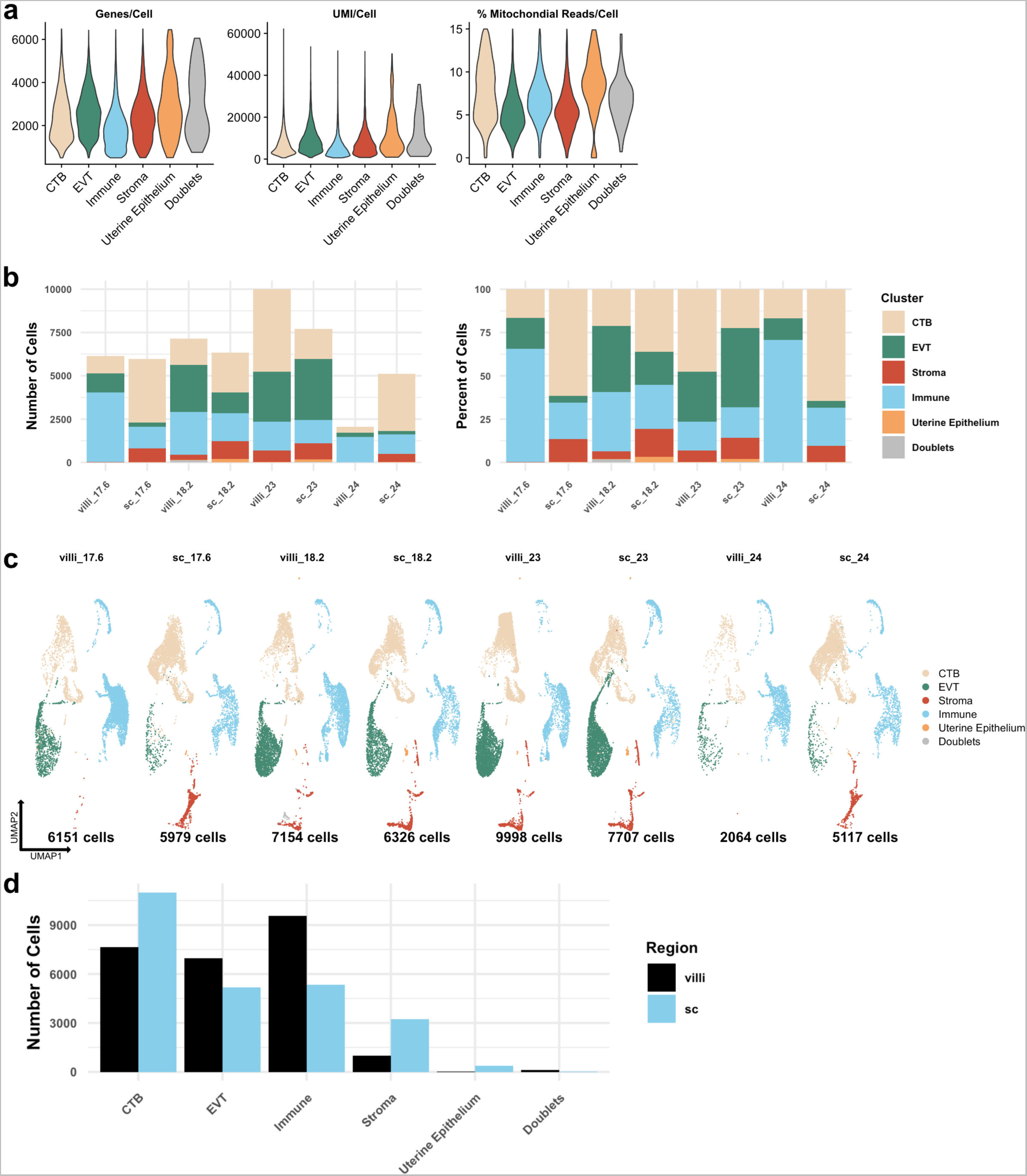
Metrics of the integrated dataset. **(a)** Violin plots of the number of unique genes (Left), number of UMI (Middle), and the percent of mitochondrial reads (Right) per cell for each broad cell type cluster. **(b)** The total number of cells (Left) and the percent of the total library (Right) in each broad cell type cluster from each placental sample are shown. **(c)** UMAP projections of the integrated dataset shown by each placental sample. Colors correspond to each broad cell type cluster in the legend at the right. The number of cells analyzed from each placental sample is listed beneath. **(d)** The number of cells in each broad cell type cluster from each placental region (VC – black; SC – blue).

**Figure 1-S2.**
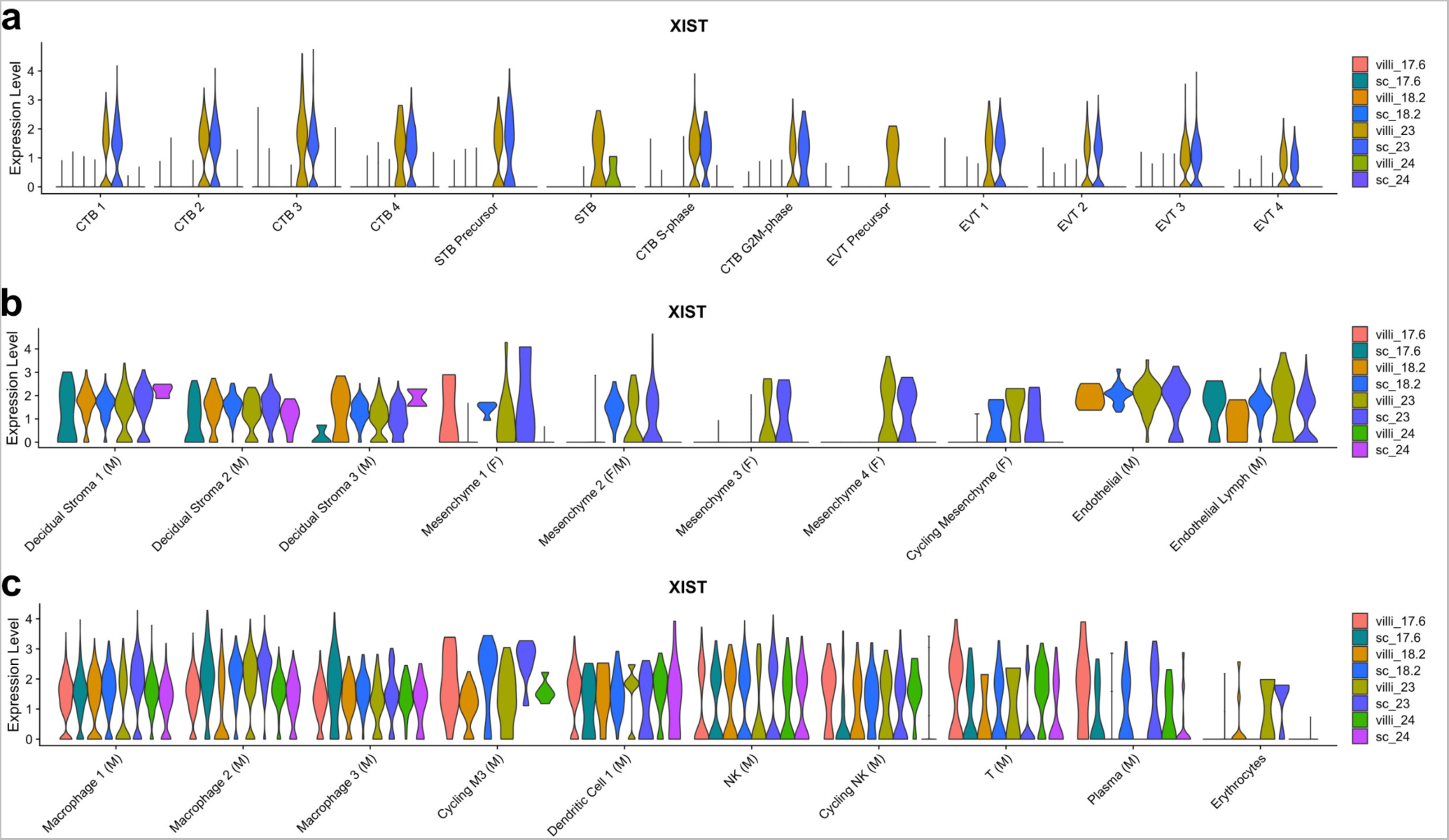
*XIST* expression by sample. **(a)** Violin plot of *XIST* transcript expression in each sample for each trophoblast cluster. Since all trophoblast are of fetal origin, samples expressing *XIST* are XX and samples with no *XIST* expression are XY. **(b)** Violin plot of *XIST* transcript expression in each sample for each stromal cluster. Clusters with XIST expression in samples other than GW23 are of maternal origin. Clusters with XIST expression only in GW23 are of fetal origin. **(c)** Violin plot of *XIST* transcript expression in each sample for each immune cluster. Clusters with XIST expression in samples other than GW23 are of maternal origin. Clusters with XIST expression only in GW23 are of fetal origin.

**Figure 1-S3.**
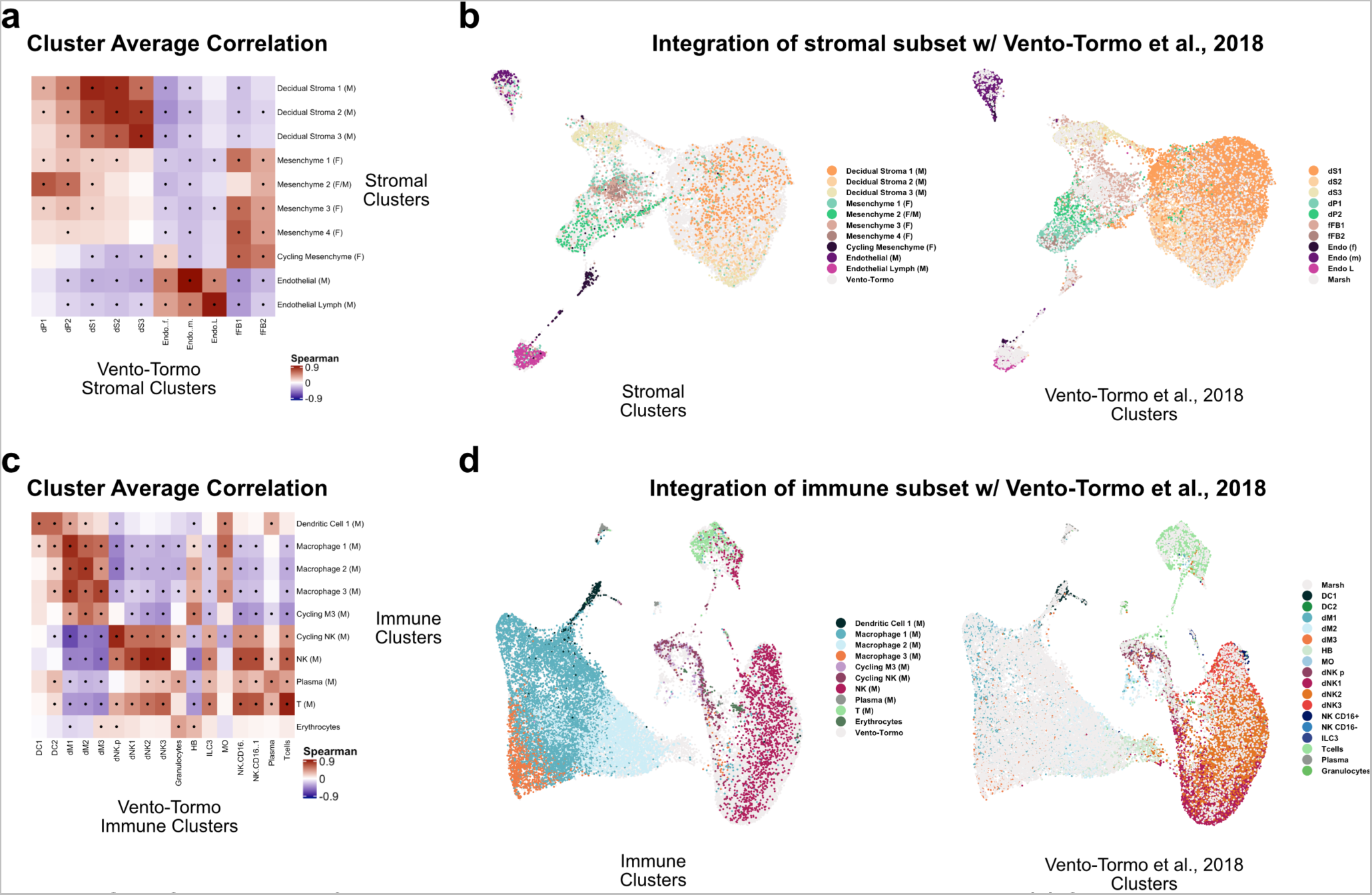
Comparison of stroma and immune clusters to Vento-Tormo et al., 2018. **(a)** Spearman correlations between the average expression within each stromal cell cluster identified in this study (y-axis) and in Vento-Tormo et al., 2018 (x-axis). Black dots denote p-value ≤ 0.05 derived from permutation testing for each correlation. **(b)** UMAP plot of the integration of stromal cells identified in this study and in Vento-Tormo et al., 2018. On the left, cells originating from this study are colored by their independently derived cluster annotations. On the right, cells originating from Vento-Tormo et al., 2018 are colored by their independently derived cluster annotations. **(c)** Spearman correlations between the average expression within each immune cell cluster identified in this study (y-axis) and in Vento-Tormo et al., 2018 (x-axis). Black dots denote p-value ≤ 0.05 derived from permutation testing for each correlation. **(d)** UMAP plot of the integration of immune cells identified in this study and in Vento-Tormo et al., 2018. On the left, cells originating from this study are colored by their independently derived cluster annotations. On the right, cells originating from Vento-Tormo et al., 2018 are colored by their independently derived cluster annotations.

**Figure 1-S4.**
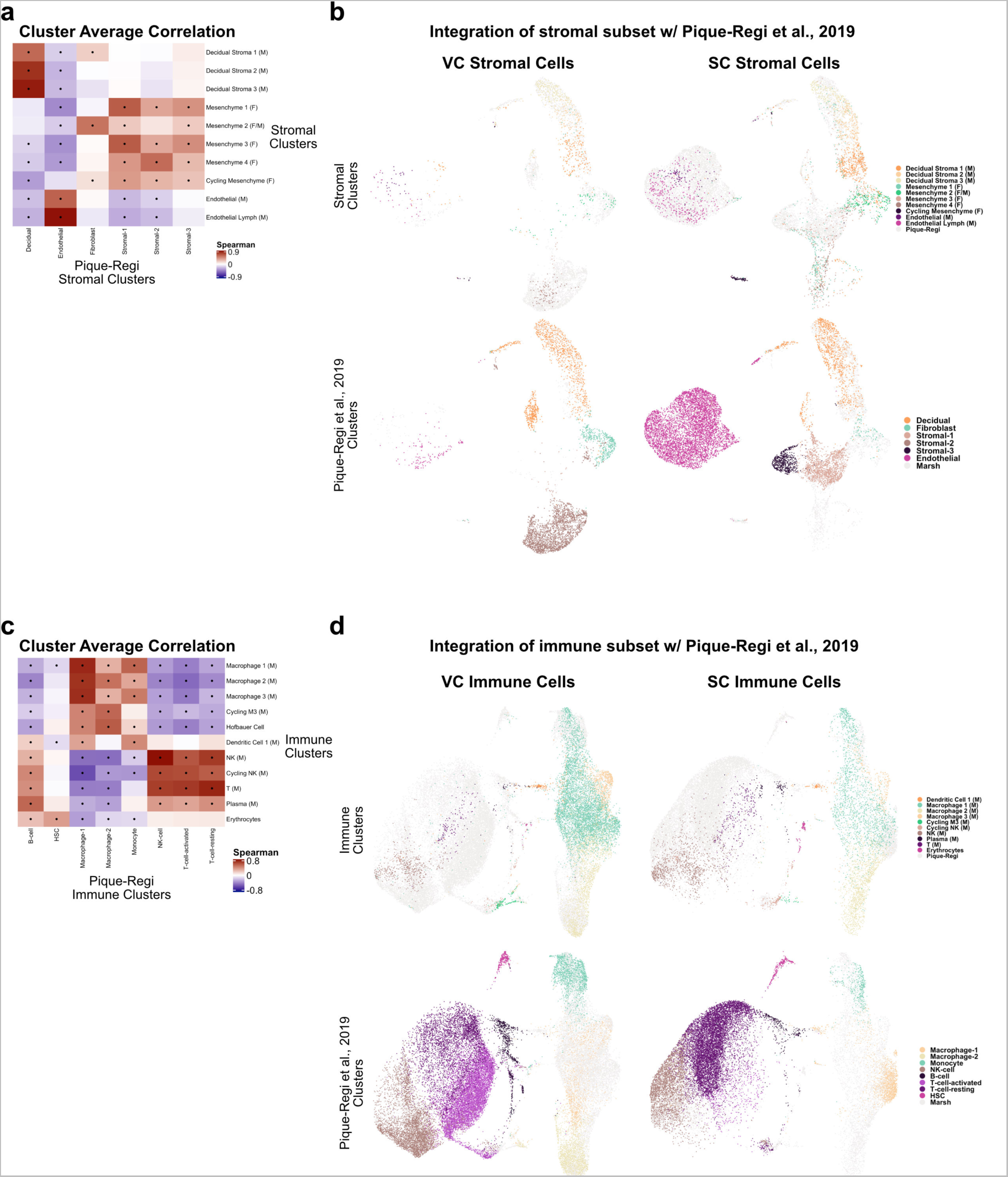
Comparison of stroma and immune clusters to Pique-Regi et al., 2019. **(a)** Spearman correlations between the average expression within each stromal cell cluster identified in this study (y-axis) and in Pique-Regi et al., 2019 (x-axis). Black dots denote p-value ≤ 0.05 derived from permutation testing for each correlation. **(b)** UMAP plots of the integration of stromal cells identified in this study and in Pique-Regi et al., 2019. Cells originating from the VC are highlighted on the left and cells originating from the SC are highlighted on the right colored by their independently derived cluster annotations. Cells originating from this study are highlighted on the top row and cells originating from Pique-Regi et al., 2019 are highlighted on the bottom row colored by their independently derived cluster annotations. **(c)** Spearman correlations between the average expression within each immune cell cluster identified in this study (y-axis) and in Pique-Regi et al., 2019 (x-axis). Black dots denote p-value ≤ 0.05 derived from permutation testing for each correlation. **(d)** UMAP plot of the integration of immune cells identified in this study and in Pique-Regi et al., 2019. Cells originating from the VC are highlighted on the left and cells originating from the SC are highlighted on the right colored by their independently derived cluster annotations. Cells originating from this study are highlighted on the top row and cells originating from Pique-Regi et al., 2019 are highlighted on the bottom row colored by their independently derived cluster annotations.

**Figure 1-S5.**
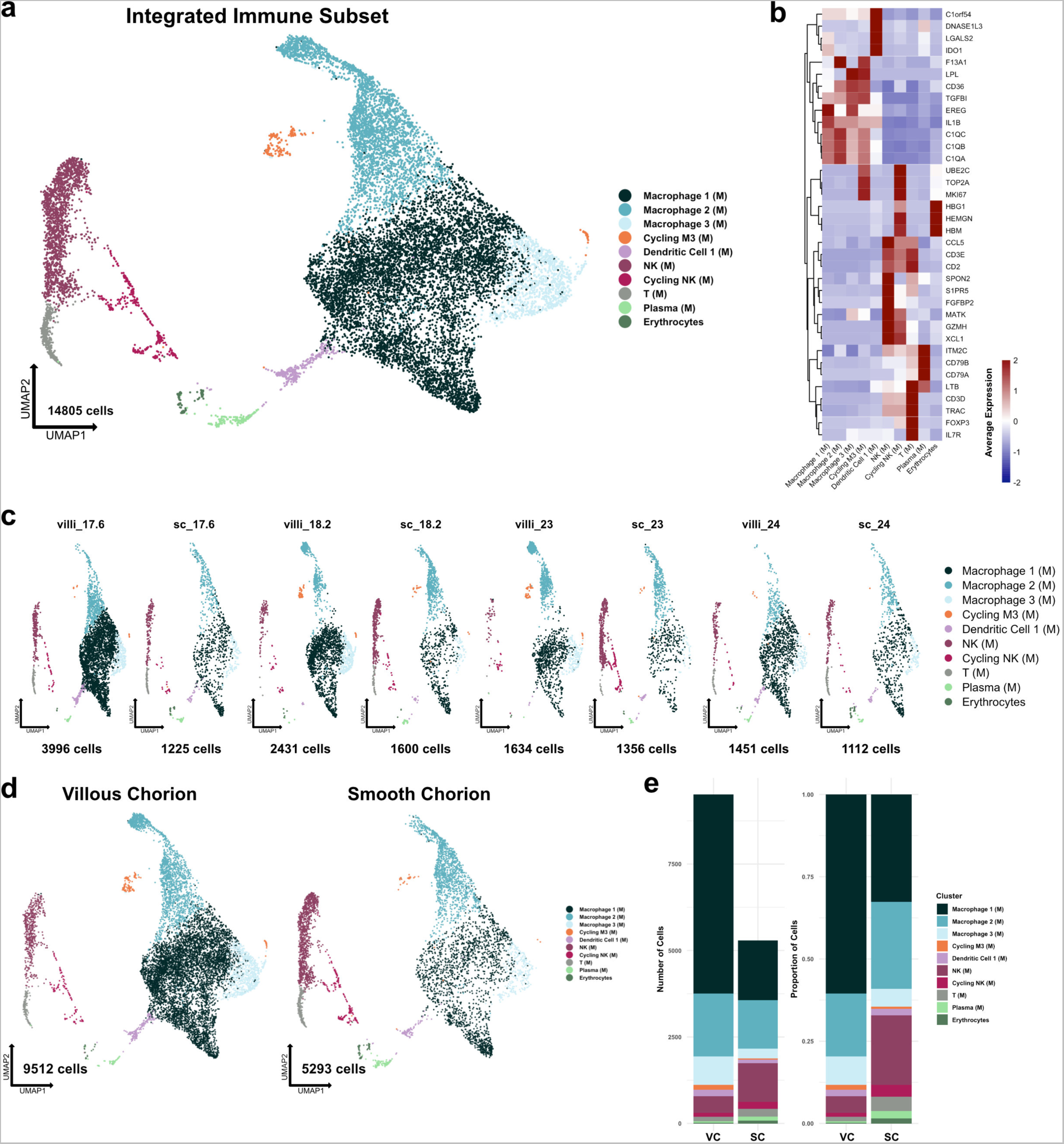
Metrics and markers of the immune cell subset. **(a)** UMAP projection of all subclustered immune cells (n=14,805). Colors correspond to the clusters in the legend at the right. **(b)** Heatmap of selected marker genes of each immune cell cluster. Expression was displayed as the scaled mean expression in the cluster. **(c)** UMAP projection of immune cells shown by sample. **(d)** UMAP projection of immune cells shown by region of origin. **(e)** Stacked bar chart of the number of cells (left) or proportion of cells (right) in each cluster by region of origin.

**Figure 1-S6.**
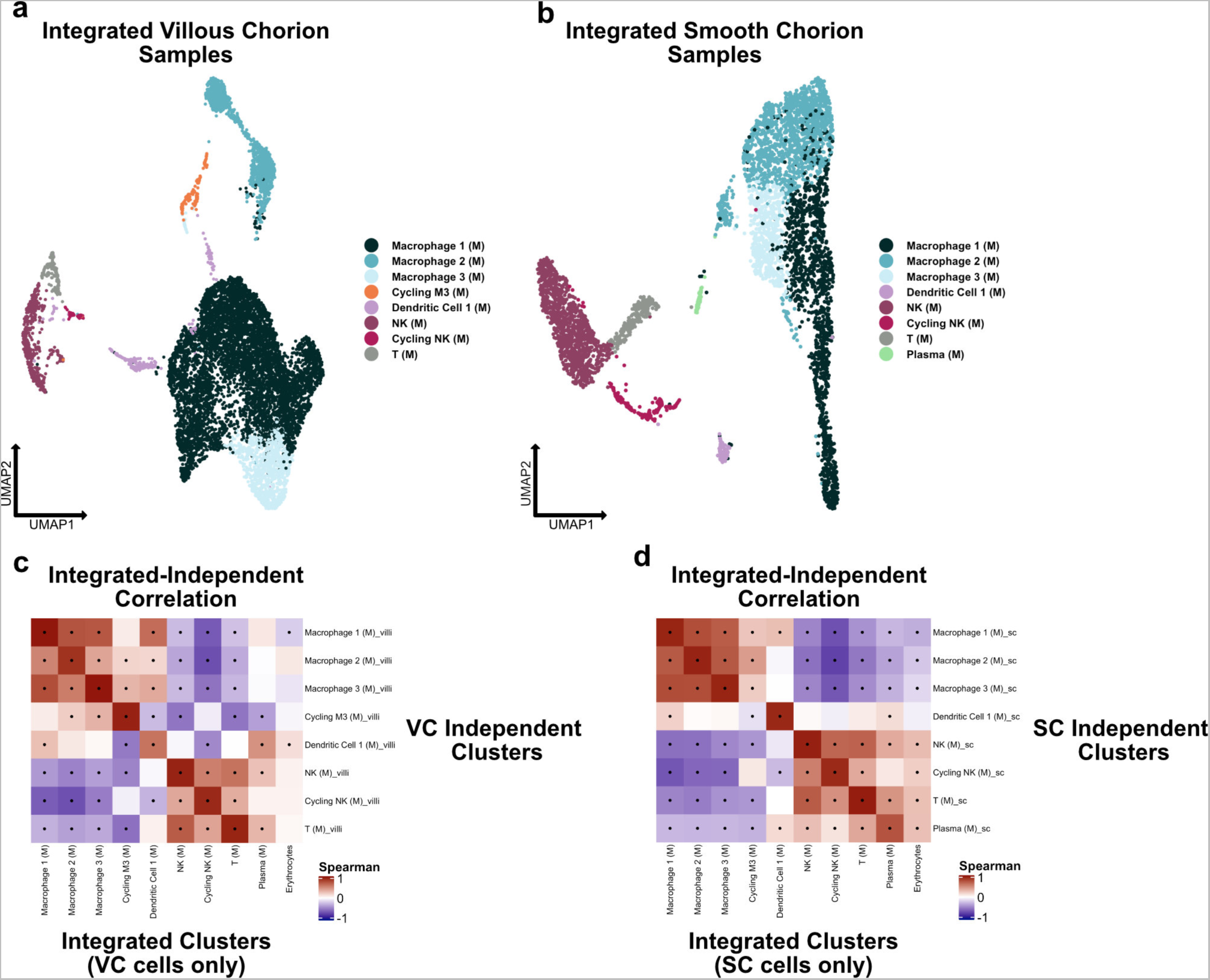
Independent analysis of VC and SC immune cells. **(a)** UMAP projection of subclustered stromal cells isolated from the VC. Colors correspond to the clusters at the right. **(b)** UMAP projection of subclustered stromal cells isolated from the SC. Colors correspond to the clusters at the right. **(c)** Spearman correlations between the average expression within independently derived VC clusters (y-axis) and the clusters from the integrated dataset (Figure 2). **(d)** Spearman correlations between the average expression within independently derived SC clusters (y-axis) and the clusters from the integrated dataset (Figure 2). Black dots denote p-value ≤ 0.05 derived from permutation testing for each correlation plot in c and d.

**Figure 1-S7.**
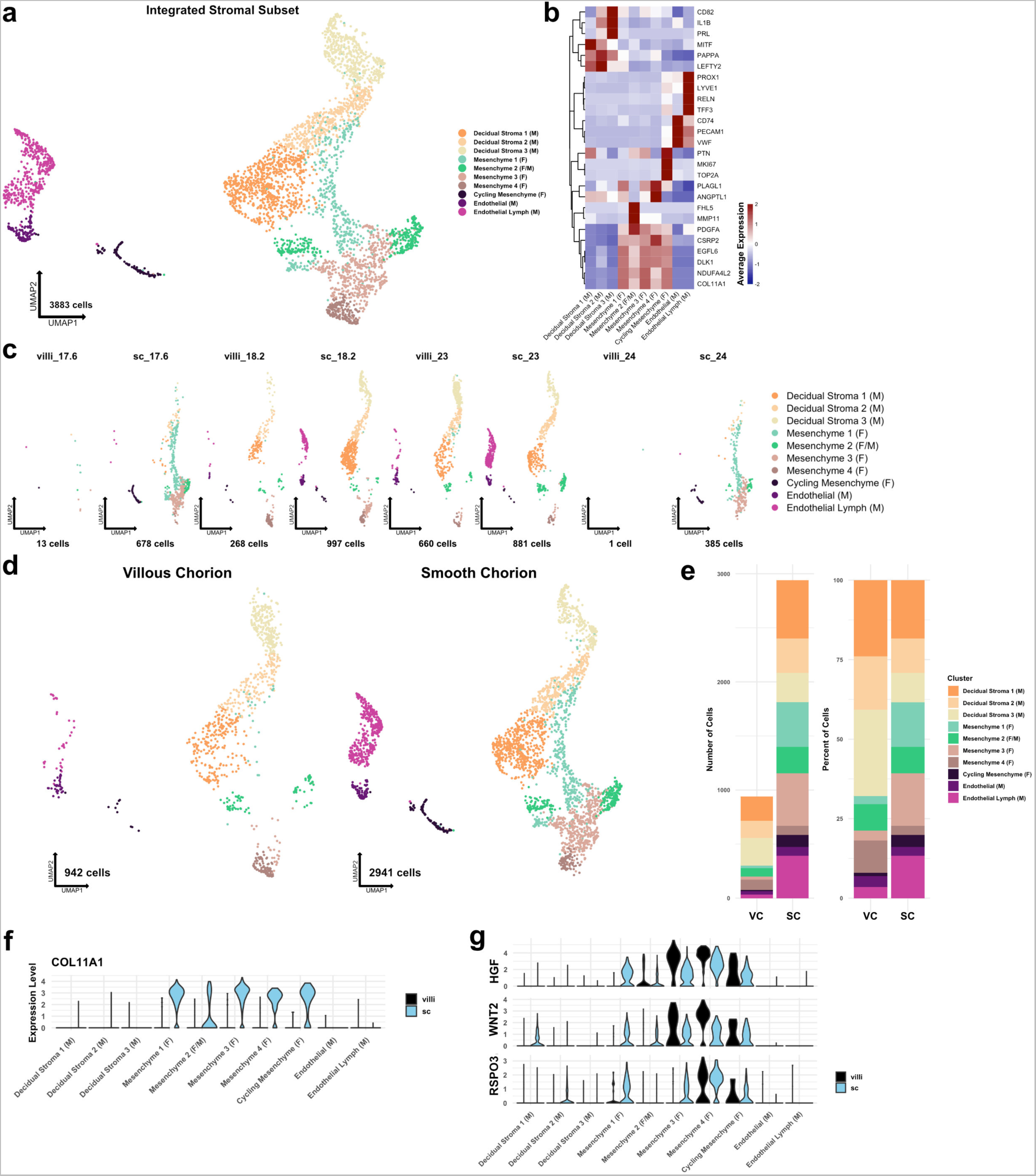
Metrics and markers of the stromal cell subset. **(a)** UMAP projection of subclustered stromal cells (n=3,883). Colors correspond to the clusters in the legend at the right. **(b)** Heatmap of selected marker genes for each stromal cell cluster. Expression was displayed as the scaled mean expression in the cluster. **(c)** UMAP projection of stromal cells shown by sample. **(d)** UMAP projection of stromal cells shown by region of origin. **(e)** Stacked bar chart of the number of cells (left) or percent of cells (right) in each cluster by region of origin. **(f)** Violin plot of COL11A1 expression in each cluster shown by each region, showing expression in only SC cells **(g)** Violin plots of HGF, WNT2, and RSPO3 expression in each cluster and shown by each region, showing expression in both VC and SC cells.

**Figure 1-S8.**
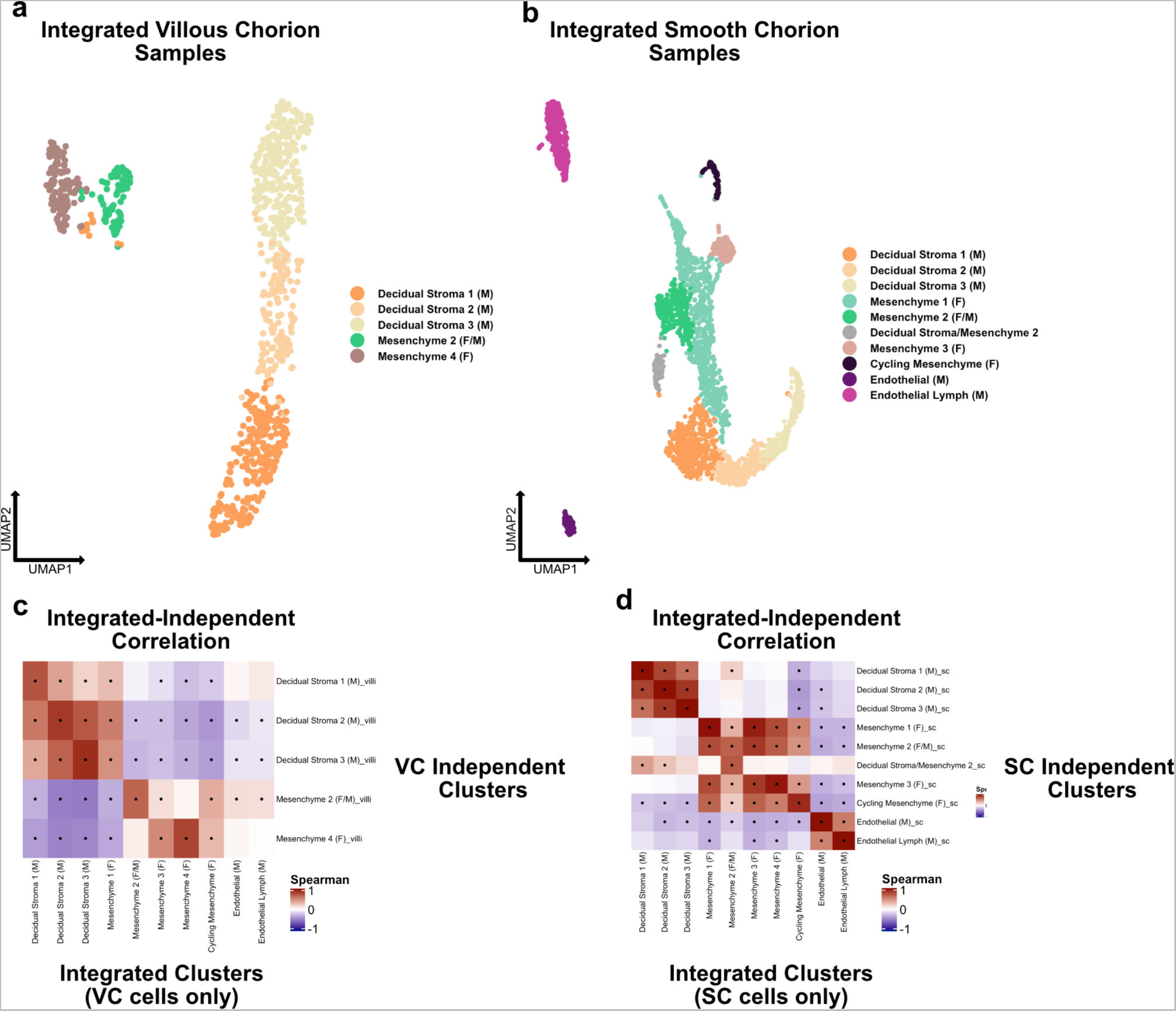
Independent analysis of VC and SC stromal cells. **(a)** UMAP projection of subclustered immune cells isolated from the VC. Colors correspond to the clusters at the right. **(b)** UMAP projection of subclustered immune cells isolated from the SC. Colors correspond to the clusters at the right. **(c)** Spearman correlations between the average expression within independently derived VC clusters (y-axis) and the clusters from the integrated dataset (Figure 2). **(d)** Spearman correlations between the average expression within independently derived SC clusters (y-axis) and the clusters from the integrated dataset (Figure 2). Black dots denote p-value ≤ 0.05 derived from permutation testing for each correlation plot in c and d.

**Figure 2-S1.**
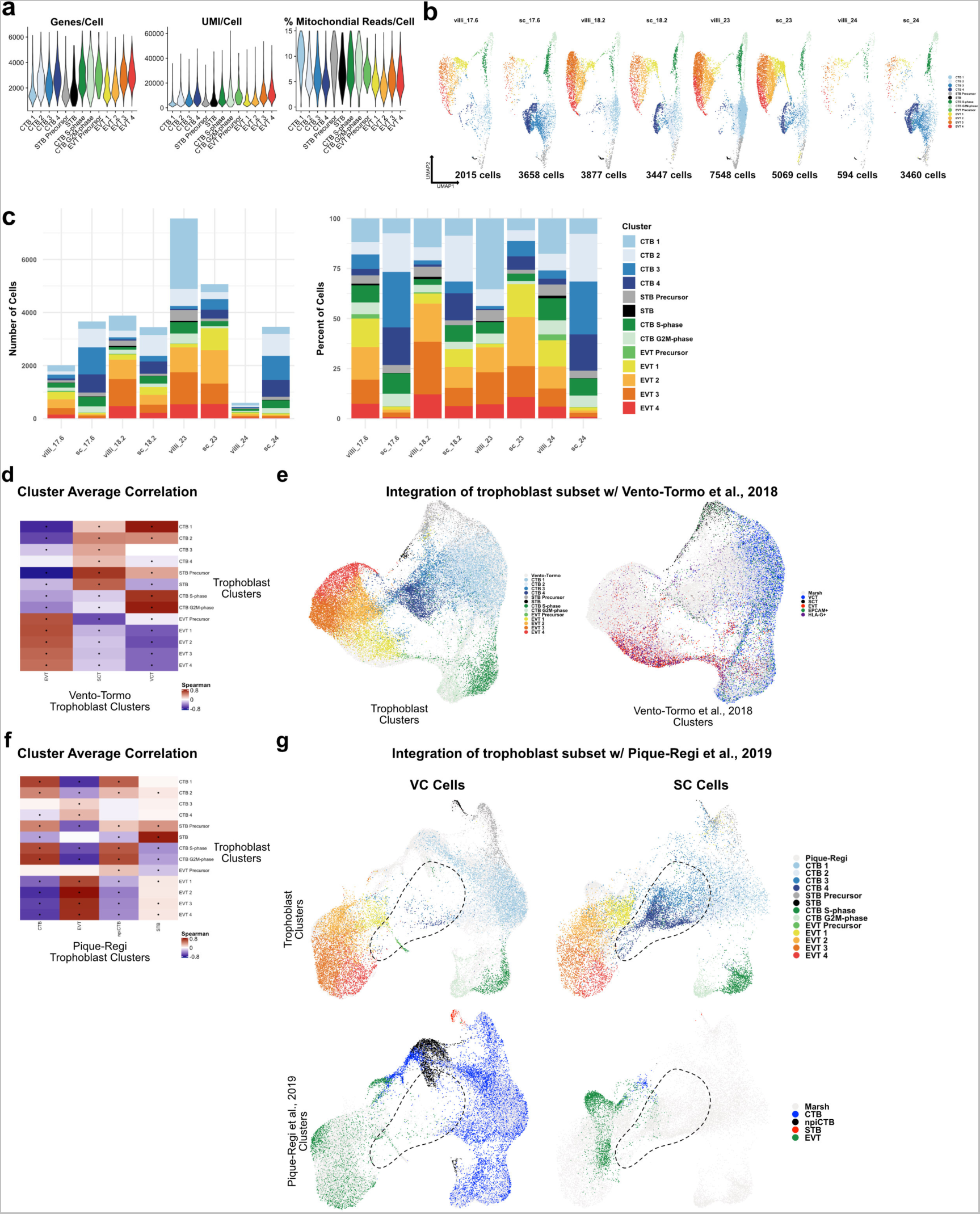
Metrics of the trophoblast subset. **(a)** Violin plots of the number of unique genes (left), number of UMI (middle), and the percent of mitochondrial reads (right) per cell for each trophoblast cluster. **(b)** UMAP projections of the trophoblast dataset shown by each placental sample. Colors correspond to each trophoblast cluster shown in the legend at the right. The number of cells analyzed from each placental sample is listed beneath. **(c)** The number of cells (left) and the percentage (right) in each trophoblast cluster from each placental sample. **(d)** Spearman correlations between the average expression within each trophoblast cell cluster identified in this study (y-axis) and in Vento-Tormo et al., 2018 (x-axis). Black dots denote p-value ≤ 0.05 derived from permutation testing for each correlation. **(e)** UMAP plot of the integration of trophoblast cells identified in this study and in Vento-Tormo et al., 2018. On the left, cells originating from this study are colored by their independently derived cluster annotations. On the right, cells originating from Vento-Tormo et al., 2018 are colored by their independently derived cluster annotations. **(f)** Spearman correlations between the average expression within each trophoblast cell cluster identified in this study (y-axis) and in Pique-Regi et al., 2019 (x-axis). Black dots denote p-value ≤ 0.05 derived from permutation testing for each correlation. **(g)** UMAP plots of the integration of trophoblast cells identified in this study and in Pique-Regi et al., 2019. Cells originating from the VC are highlighted on the left and cells originating from the SC are highlighted on the right colored by their independently derived cluster annotations. Cells originating from this study are highlighted on the top row and cells originating from Pique-Regi et al., 2019 are highlighted on the bottom row colored by their independently derived cluster annotations. The dashed line annotates the contribution of CTB 3 and CTB 4 identified in this study, and their absence in the VC and in the Pique-Regi et al., 2019 dataset.

**Figure 2-S2.**
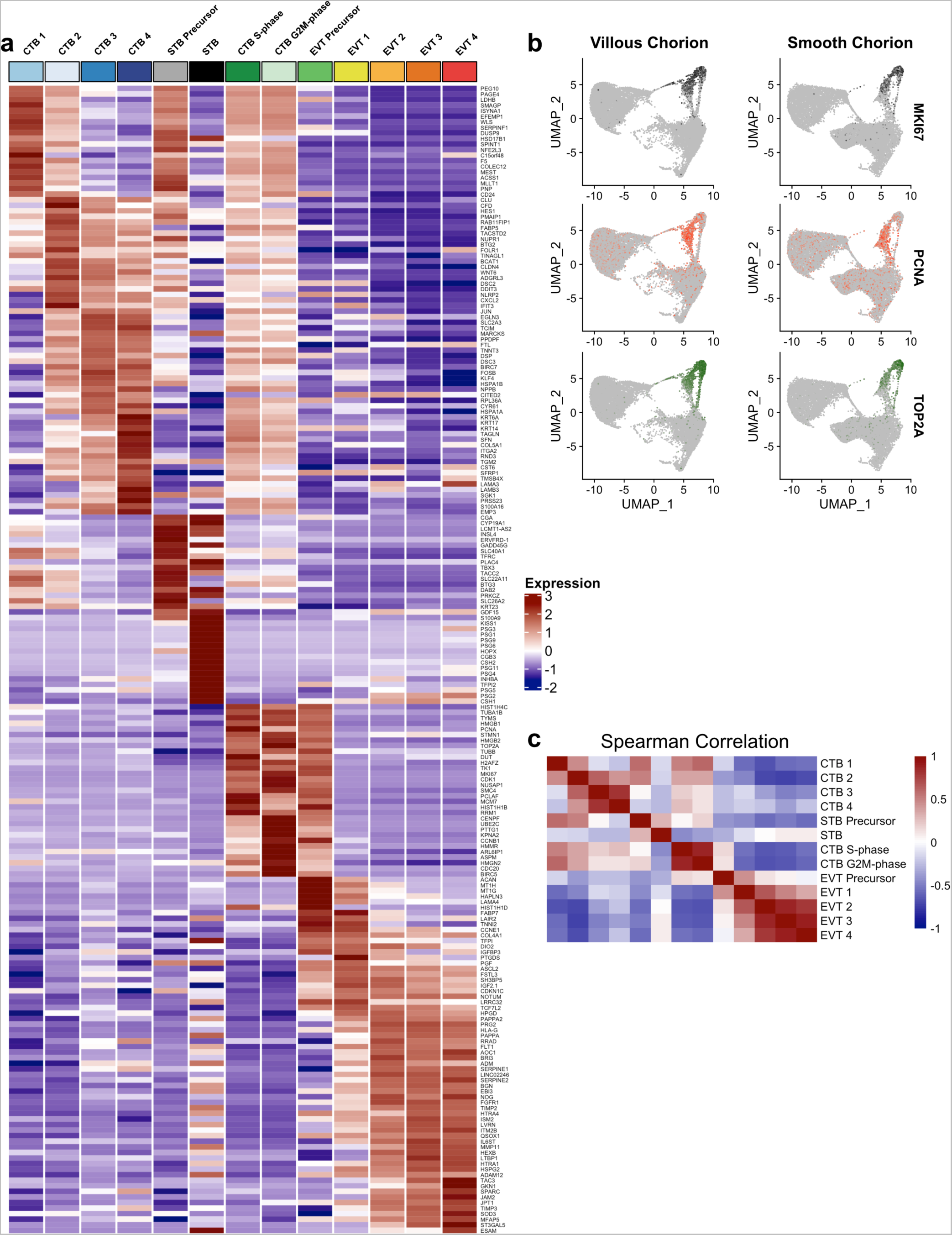
Markers of the trophoblast subset. **(a)** Heatmap of the expression of the top 20 marker genes for each trophoblast population. **(b)** Expression of phasic transcripts MKI67 (top), PCNA (middle), and TOP2A (bottom) were projected in UMAP space. **(c)** Heatmap of the Spearman Correlation coefficients between each trophoblast population.

**Figure 2-S3.**
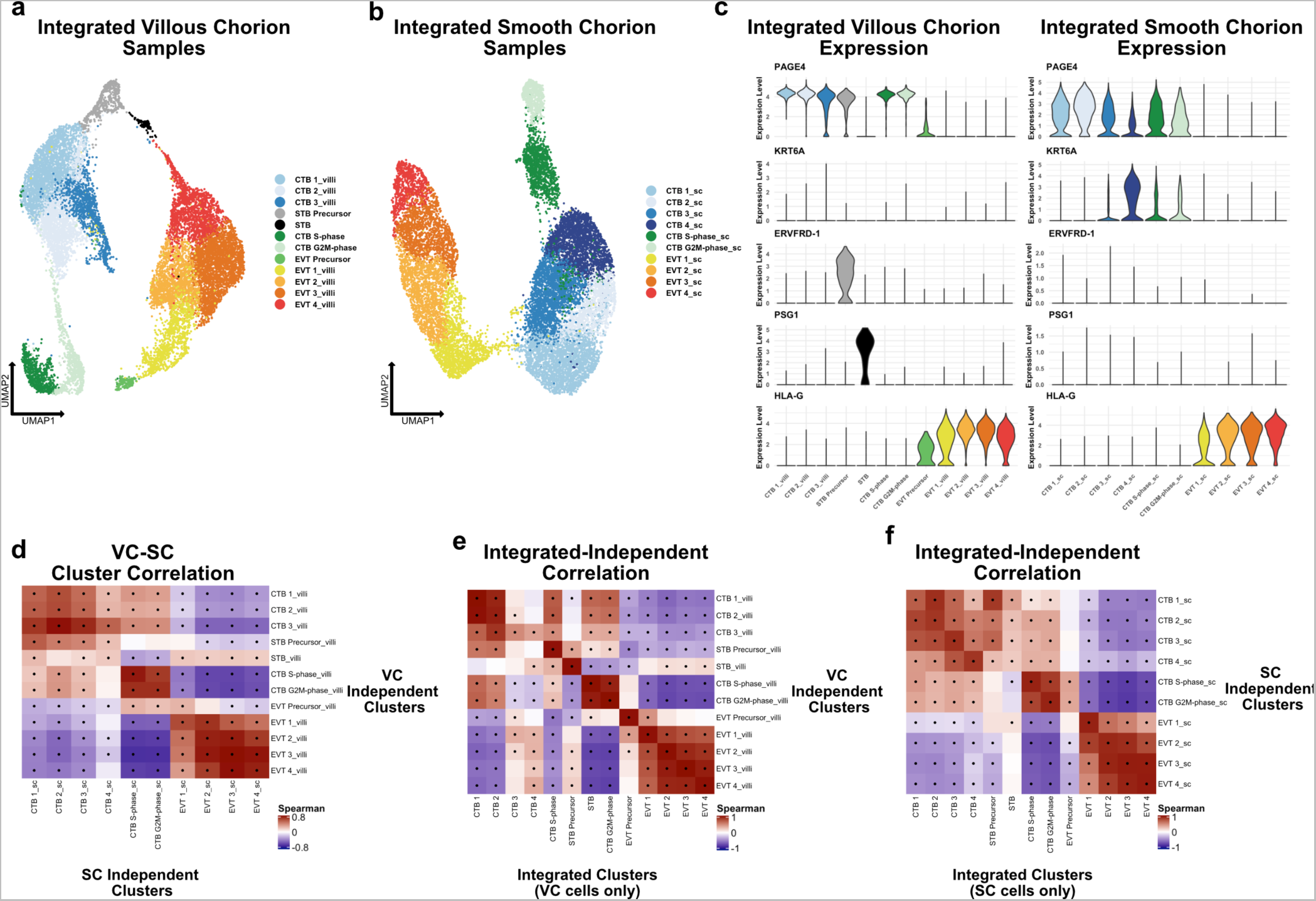
Independent analysis of VC and SC trophoblast. **(a)** UMAP projection of subclustered trophoblasts isolated from the VC. Colors correspond to the clusters at the right. **(b)** UMAP projection of subclustered trophoblasts isolated from the SC. Colors correspond to the clusters at the right. **(c)** Violin plots of transcript expression of CTB 1 marker *PAGE4*, CTB 4 marker *KRT6A*, STB Precursor marker *ERVFRD-1*, STB marker *PSG1*, and EVT marker *HLA-G* in the independently derived VC cell clusters (left) and SC cell clusters (right). **(d)** Spearman correlations between the average expression within independently derived VC clusters (y-axis) and the independently derived SC clusters (x-axis). Black dots denote p-value ≤ 0.05 derived from permutation testing for each correlation. **(e)** Spearman correlations between the average expression within independently derived VC clusters (y-axis) and the clusters from the integrated dataset (Figure 2). Black dots denote p-value ≤ 0.05 derived from permutation testing for each correlation. **(f)** Spearman correlations between the average expression within independently derived SC clusters (y-axis) and the clusters from the integrated dataset (Figure 2). Black dots denote p-value ≤ 0.05 derived from permutation testing for each correlation.

**Figure 2-S4.**
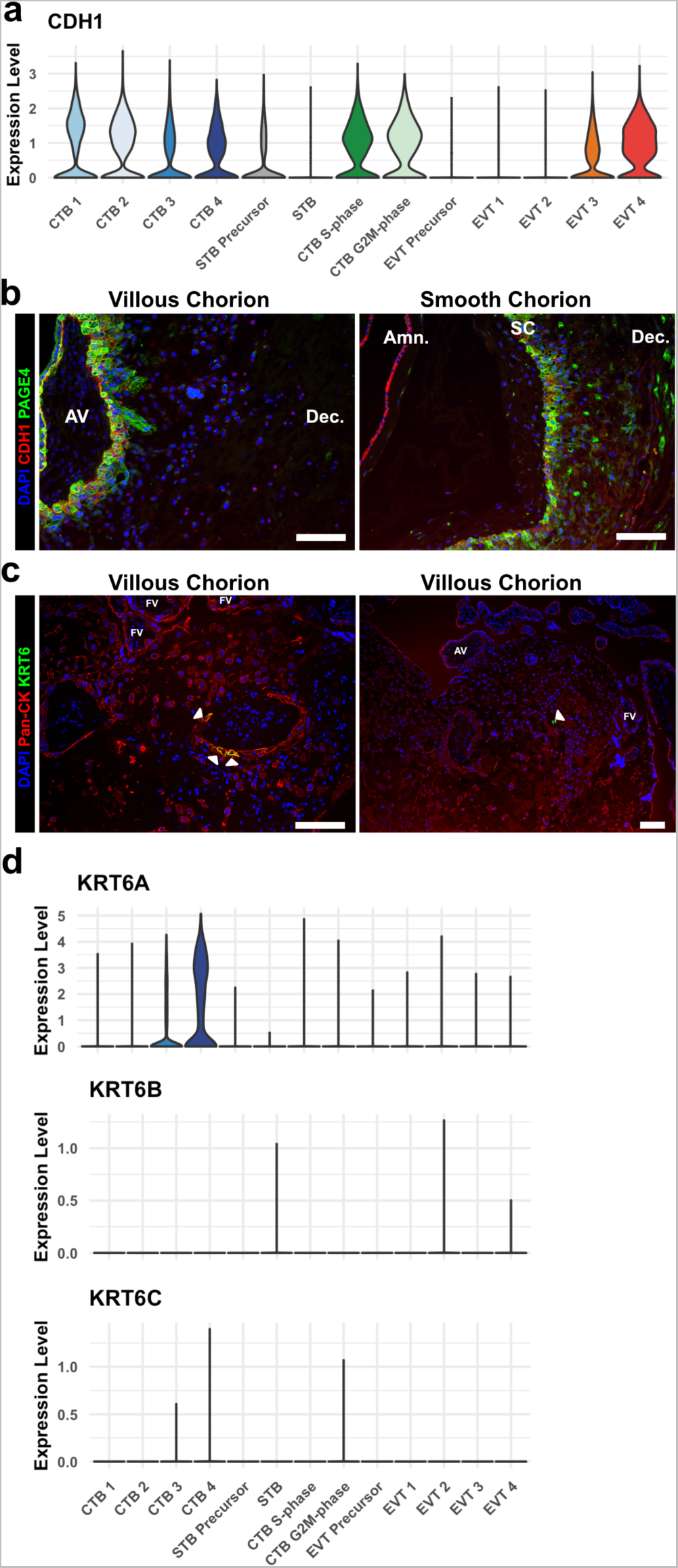
KRT6 expression in the VC region. **(a)** Violin plot of *CDH1* transcript expression across all trophoblast clusters. **(b)** Immunofluorescence co-localization of CDH1 and PAGE4 in the VC (Left) and SC (Right). **(c)** Immunofluorescence co-localization of Pan-CK (marker of all trophoblast) and KRT6 in the VC. Arrowheads denote KRT6+ cells. **(d)** Violin plots of expression of each KRT6 isoform, showing expression of only KRT6A in trophoblasts. For all images nuclei were visualized by DAPI stain; Scale bar = 100μm. Abbreviations: AV = Anchoring Villi; Amn. = Amnion; SC = Smooth Chorion epithelium; Dec. = Decidua; FV = Floating Villi.

**Figure 3-S1.**
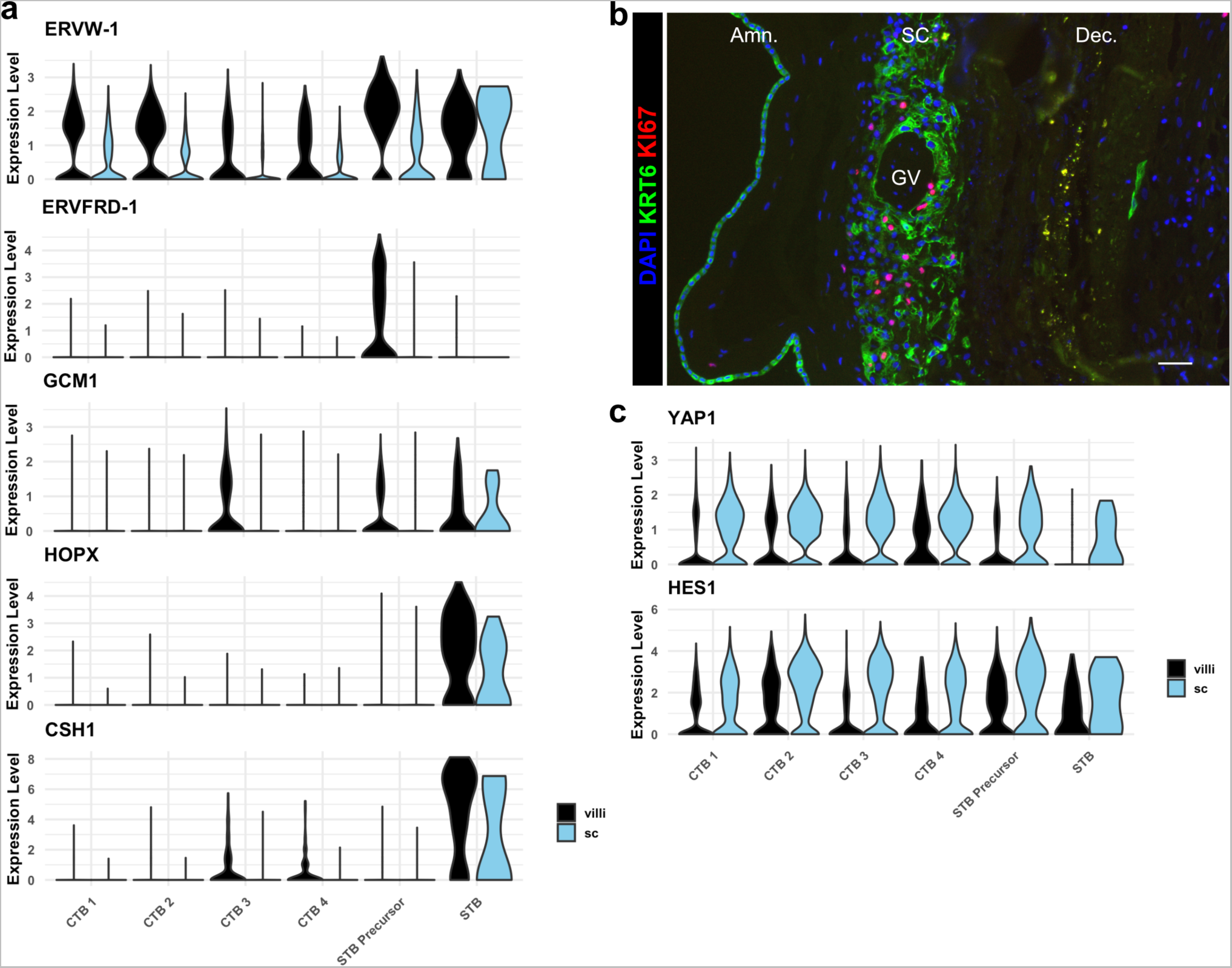
SC trophoblast display reduced expression of STB, increased expression of epithelial TFs, and proliferate. **(a)** Violin plots of expression of markers of STB differentiation in each trophoblast cluster shown by region, demonstrating reduced expression all markers in STB Precursor and STB clusters. **(c)** Violin plots of expression of YAP1 and HES1, demonstrating increasing expression in CTB1-4 and greater expression in the SC compared to the VC.

**Figure 3-S2.**
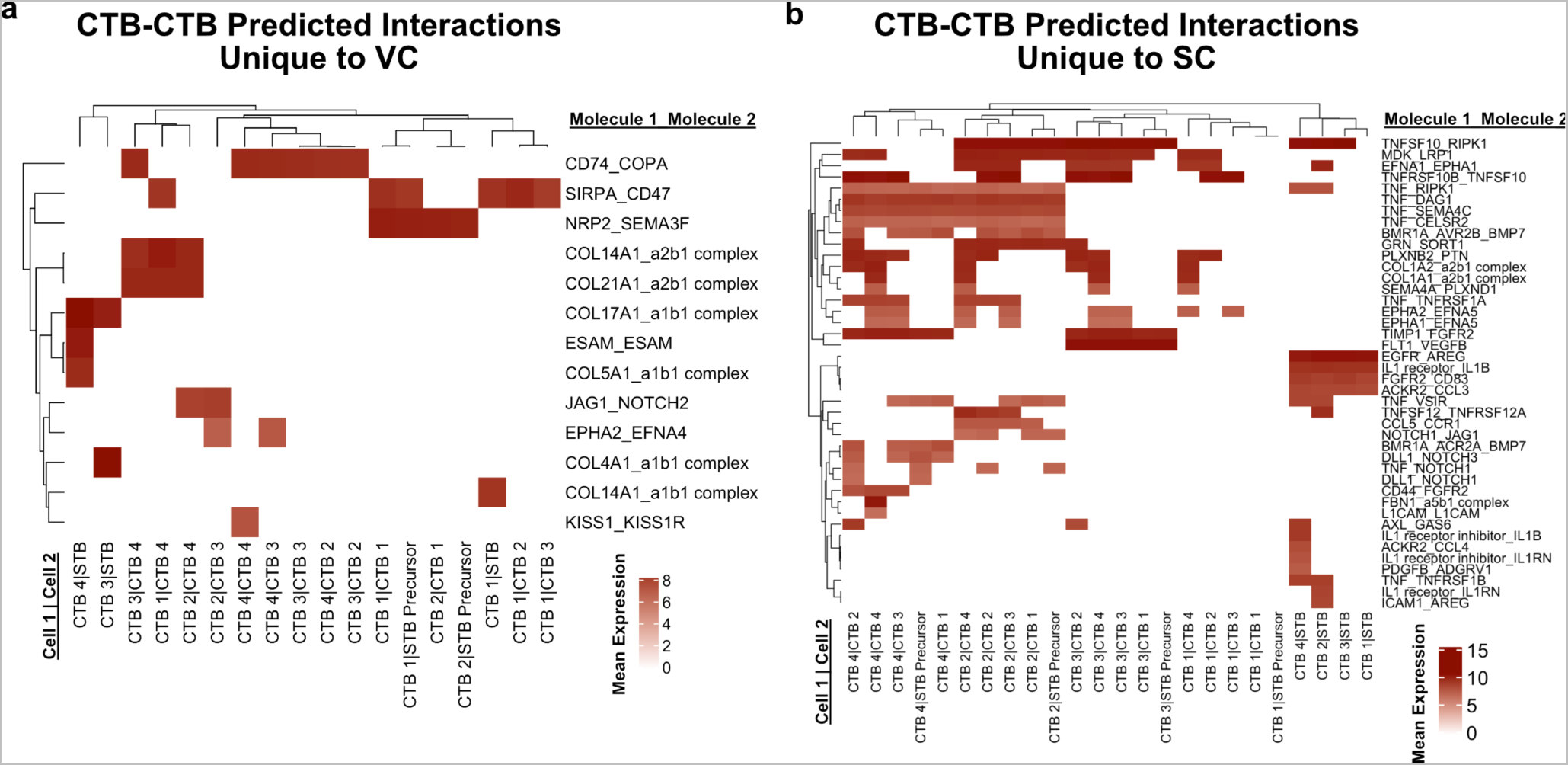
Predicted interactions between CTBs. **(a)** Predicted receptor-ligand interactions from CellPhoneDB between CTB clusters of the VC for interactions which are unique to the VC. **(b)** Predicted receptor-ligand interactions from CellPhoneDB between CTB clusters of the SC for interactions which are unique to the SC. The strength of interaction is estimated by mean expression and are plotted in the heatmaps. Receptor-ligand interactions and cell pairs are listed such that Molecule 1 is expressed by Cell 1 and Molecule 2 is expressed by Cell 2.

**Figure 4-S1.**
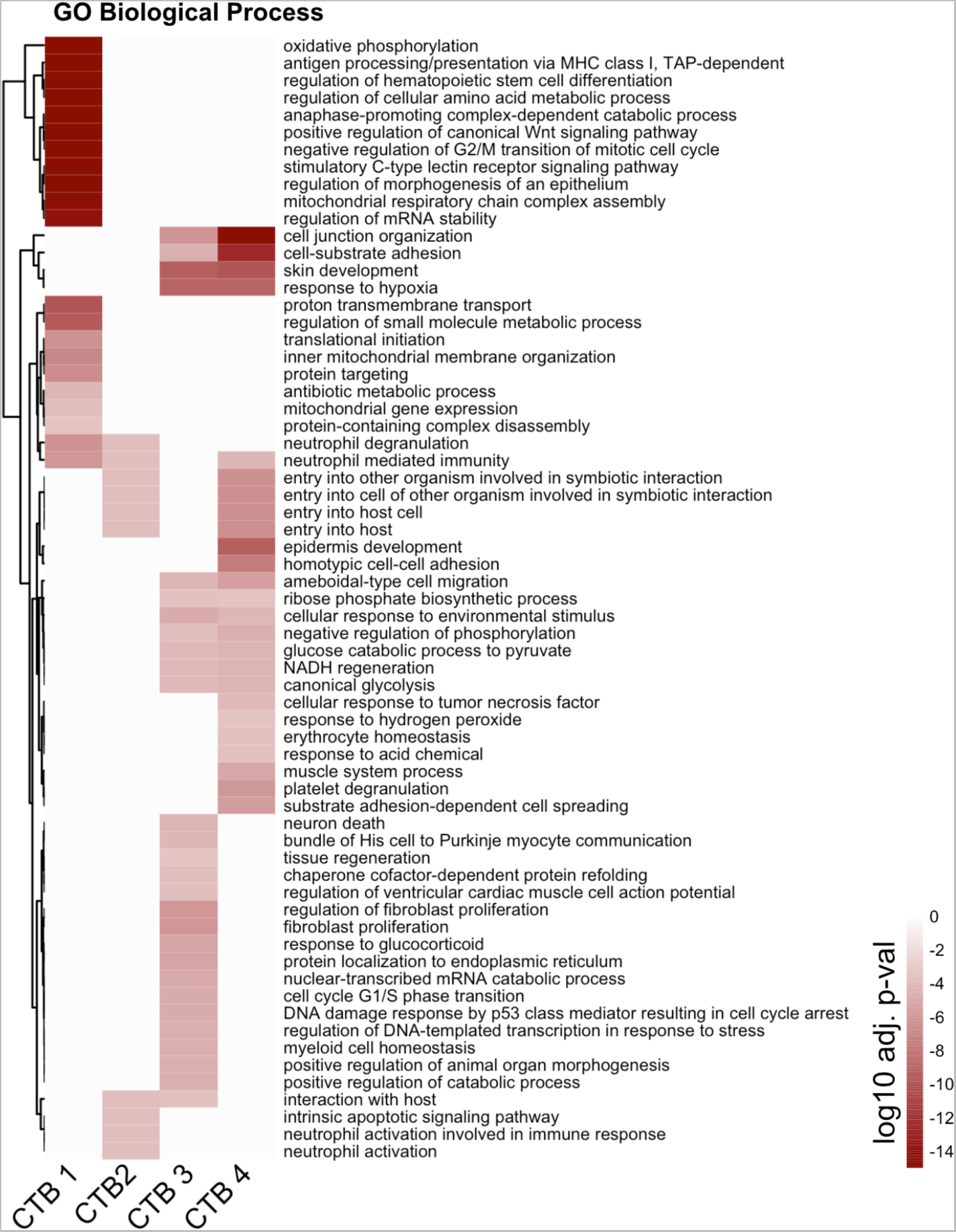
Complete CTB Gene Ontology Analysis. The unabridged Gene Ontology results from each CTB cluster, displayed as adjusted p-values. Dark red corresponds to the lowest p-values and white represents p-values greater than 0.0005. Ontology categories are organized by hierarchical clustering along the y-axis. Marker genes for each cluster were used as inputs for the analysis.

**Figure 4-S2.**
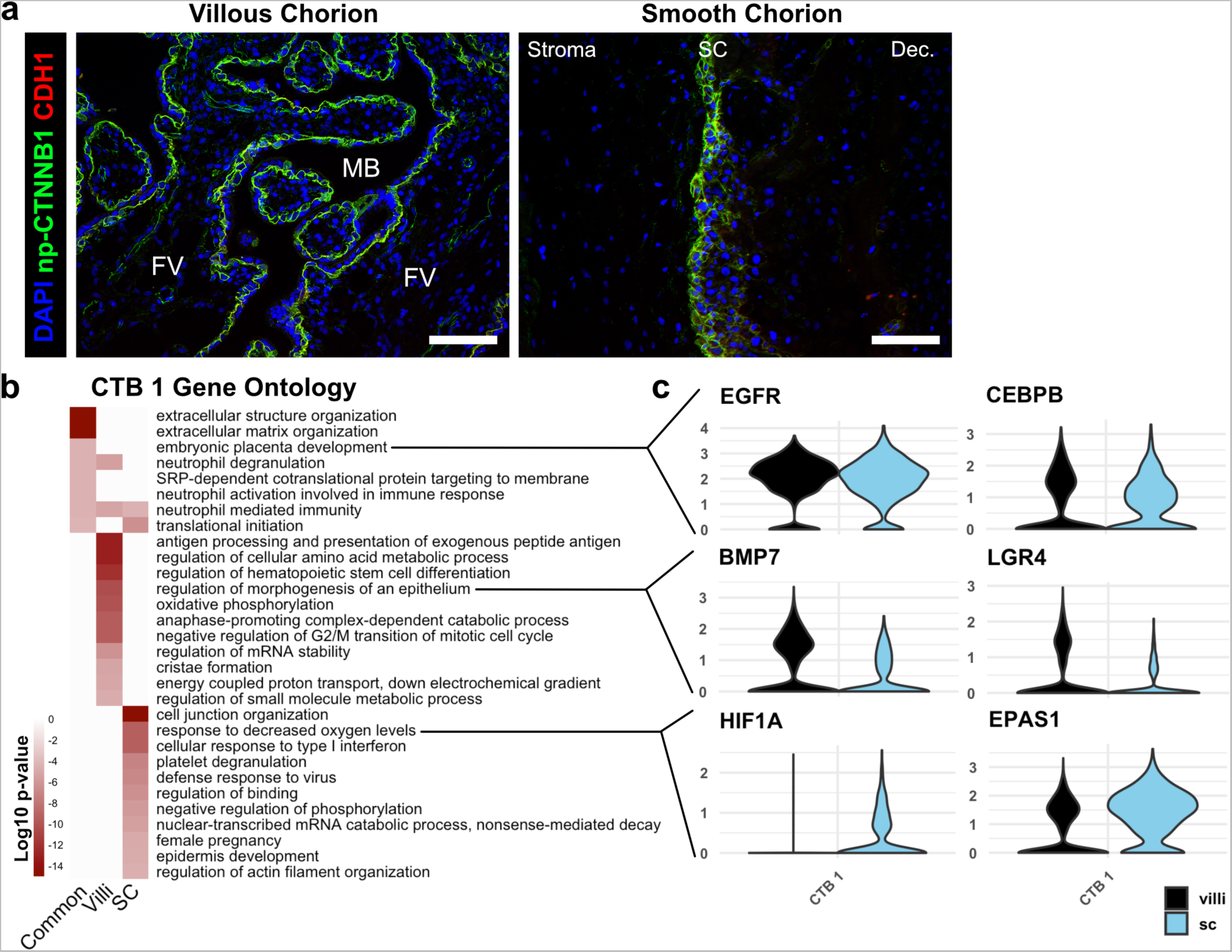
Similarities and differences between CTB 1 in VC and SC. **(a)** Immunofluorescence co-localization of CDH1 and non-phosphorylated CTNNB1 in the VC (left) and SC (right), showing identical domains of expression in both regions. For all images nuclei were visualized by DAPI stain; Scale bar = 100μm. Abbreviations: FV = Floating Villi; MB = Maternal Blood space; SC = Smooth Chorion epithelium; Dec. = Decidua. **(b)** Gene ontology analysis for CTB 1 genes common to VC and SC, genes enriched in VC, and genes enriched in SC. **(c)** Violin plots of genes within the gene ontology categories indicated in (b) for common categories (top), VC enriched categories (middle), and SC enriched categories (bottom).

**Figure 4-S3.**
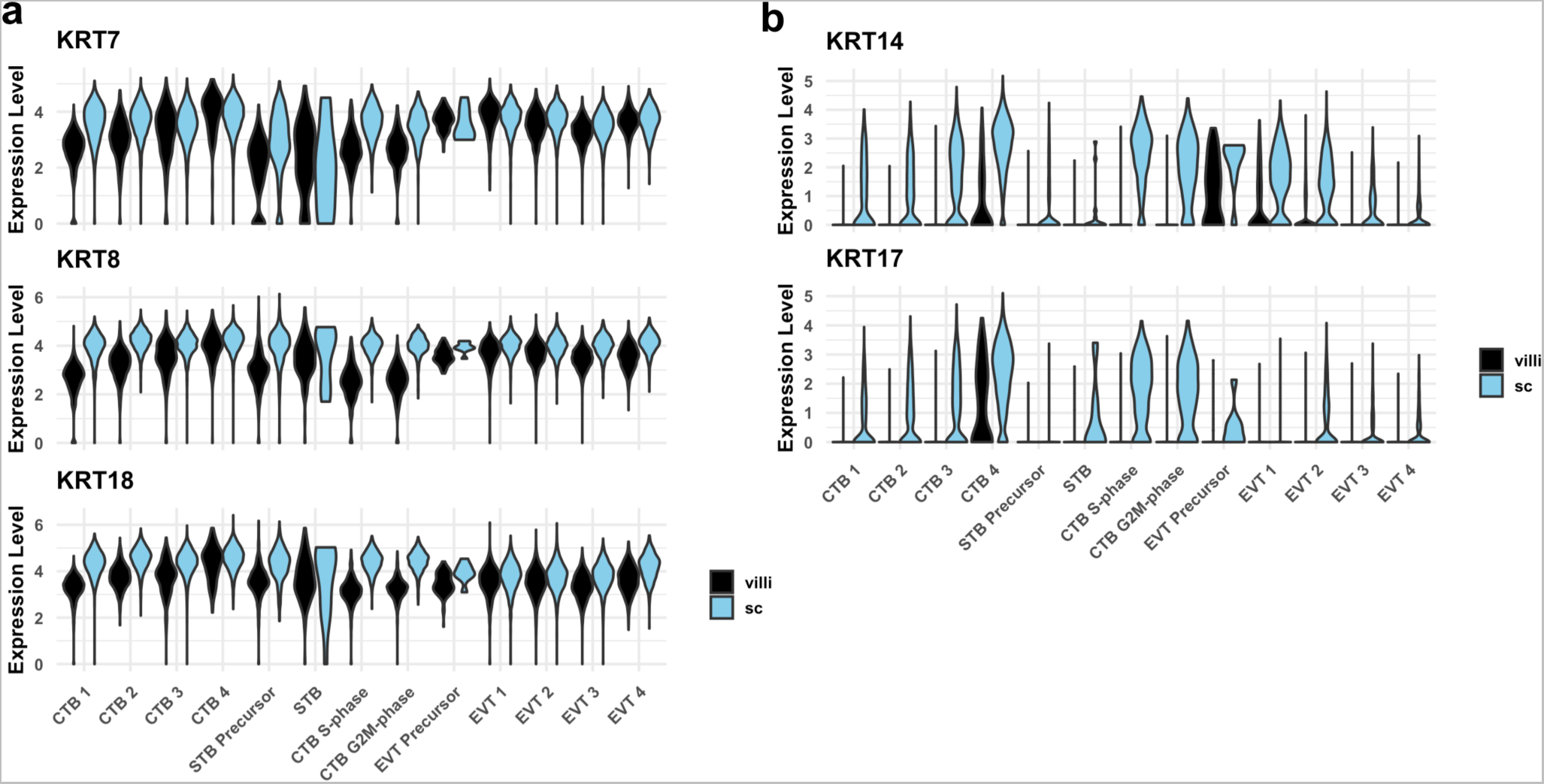
Cytokeratin expression in VC and SC trophoblast. **(a)** Violin plots of cytokeratins expressed in trophoblast in both VC and SC regions by cluster and region of origin. **(b)** Violin plots of cytokeratins expressed in SC trophoblast by cluster and region of origin.

**Figure 4-S4.**
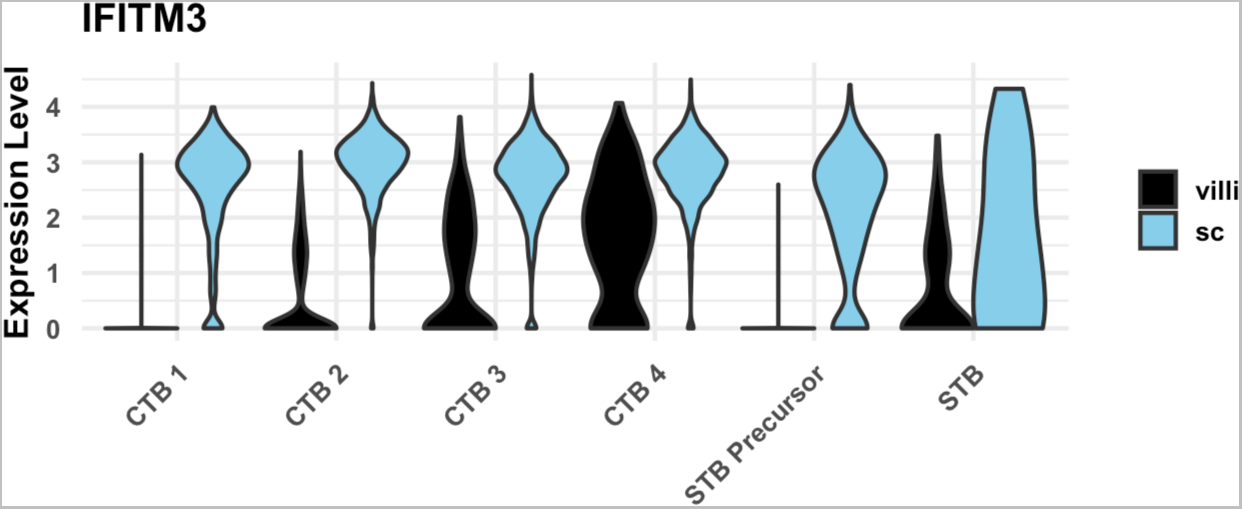
IFITM3 expression in CTB populations. **(a)**Violin plot of IFITM3 expression in trophoblast clusters by region, showing increased expression in SC compared to VC.

**Figure 5-S1.**
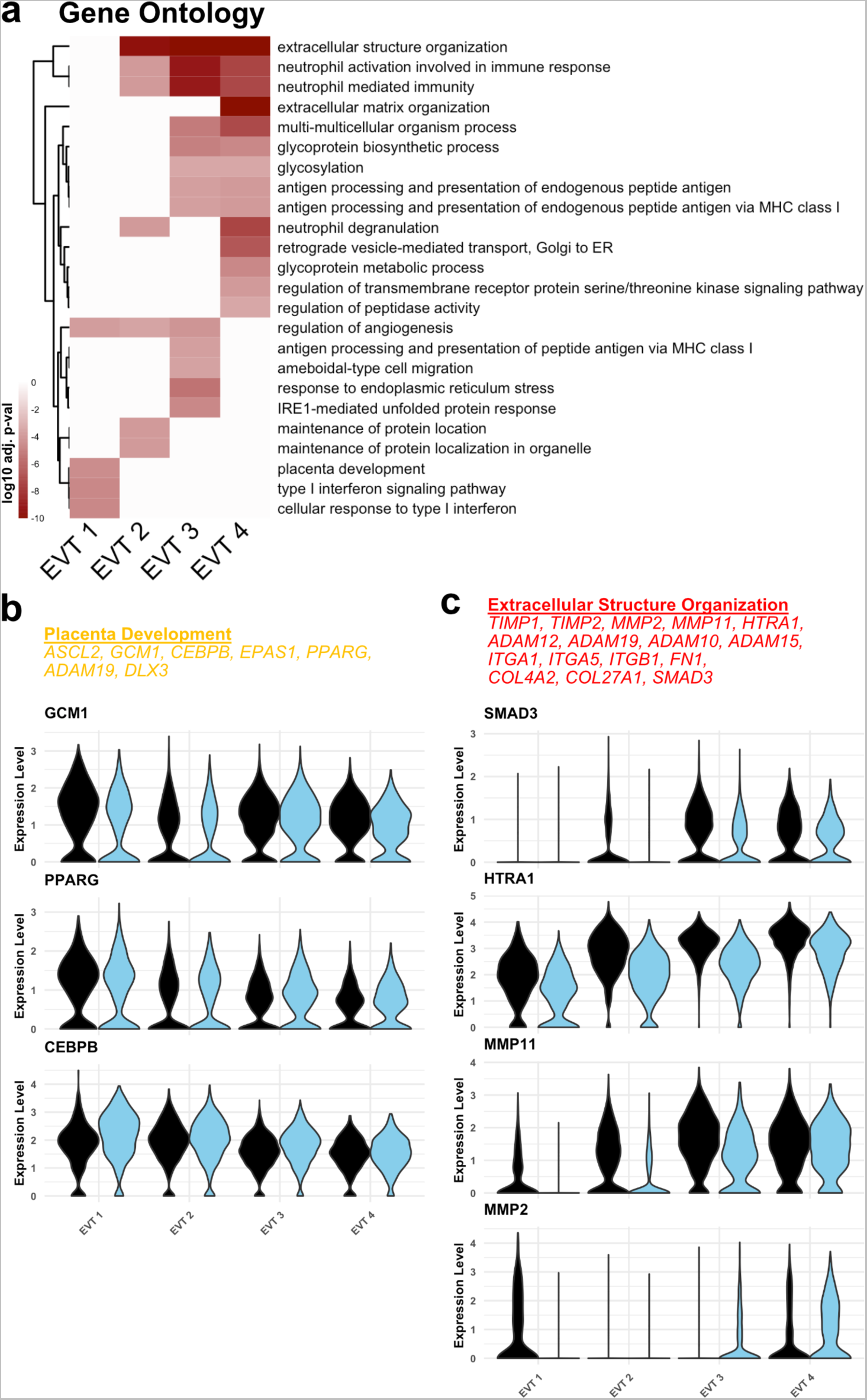
Functional annotation of EVT clusters. **(a)** Gene Ontology results from each EVT cluster displayed as adjusted p-values. Dark red corresponds to the lowest p-values and white represents p-values greater than 0.0005. Ontology categories are organized by hierarchical clustering along the y-axis. Marker genes for each cluster were used as inputs for the analysis. **(b)** Selected genes in the ‘placenta development’ gene ontology category (top). Violin plots of expression for selected genes in each EVT cluster by region. **(c)** Selected genes in the ‘Extracellular Structure Organization’ gene ontology category (top). Violin plots of expression for selected genes in each EVT cluster by region.

**Figure 5-S2.**
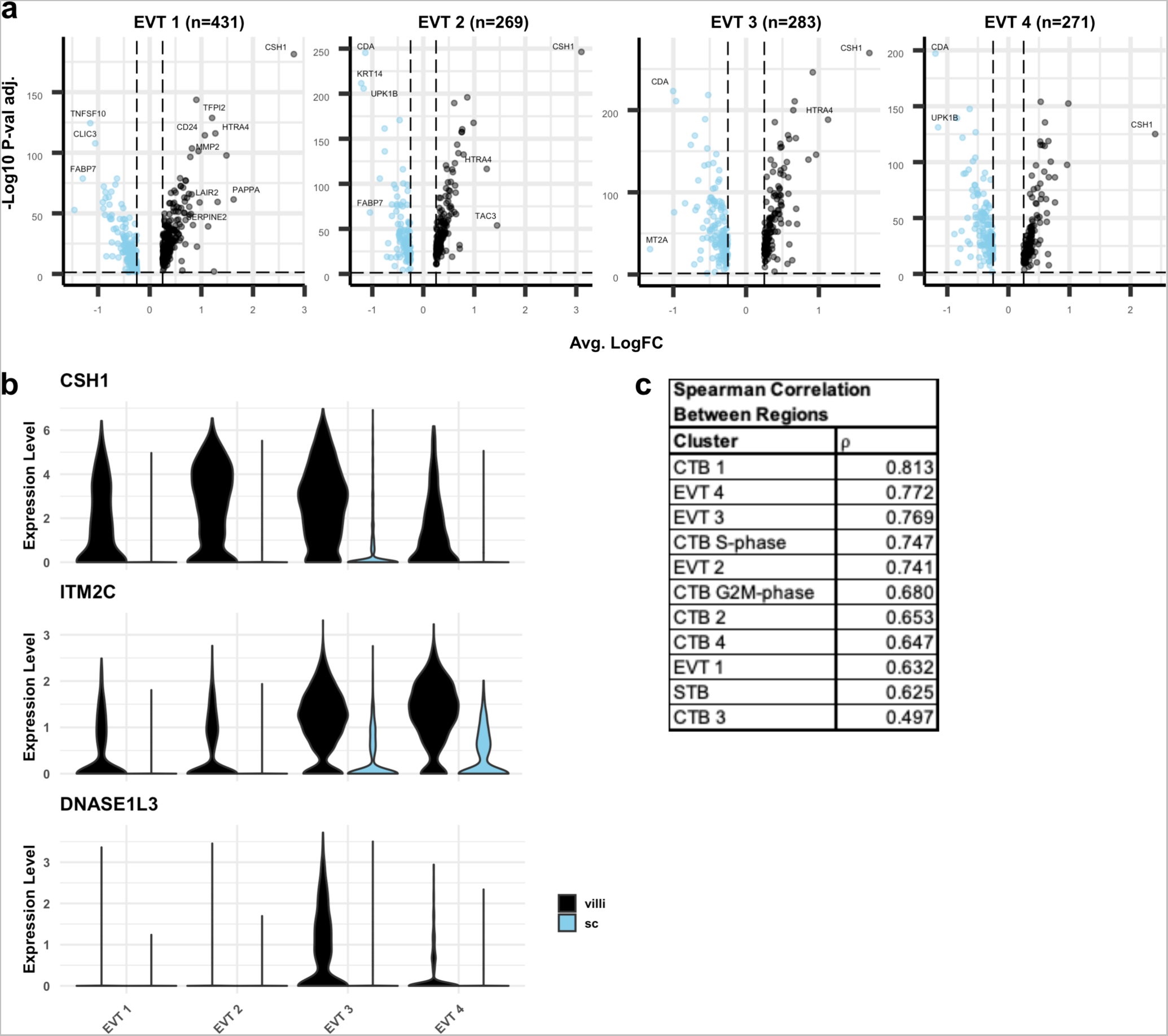
Differential expression between EVT from VC and SC. **(a)** Volcano plots showing the differentially expressed genes the VC or SC for each EVT cluster. Only significantly differentially expressed genes are plotted (p-value <0.05). Genes with higher expression in VC compared to SC are shown in black. Genes with higher expression in SC compared to VC are shown in blue. Selected differentially expressed genes are labelled. **(b)** Violin plots of VC-specific genes in EVTs. **(c)** Spearman correlation coefficients between VC and SC for each cluster ranked from greatest to least similarity across regions. STB and EVT Precursor populations are excluded due to the low number of cells recovered in the SC.

**Figure 5-S3.**
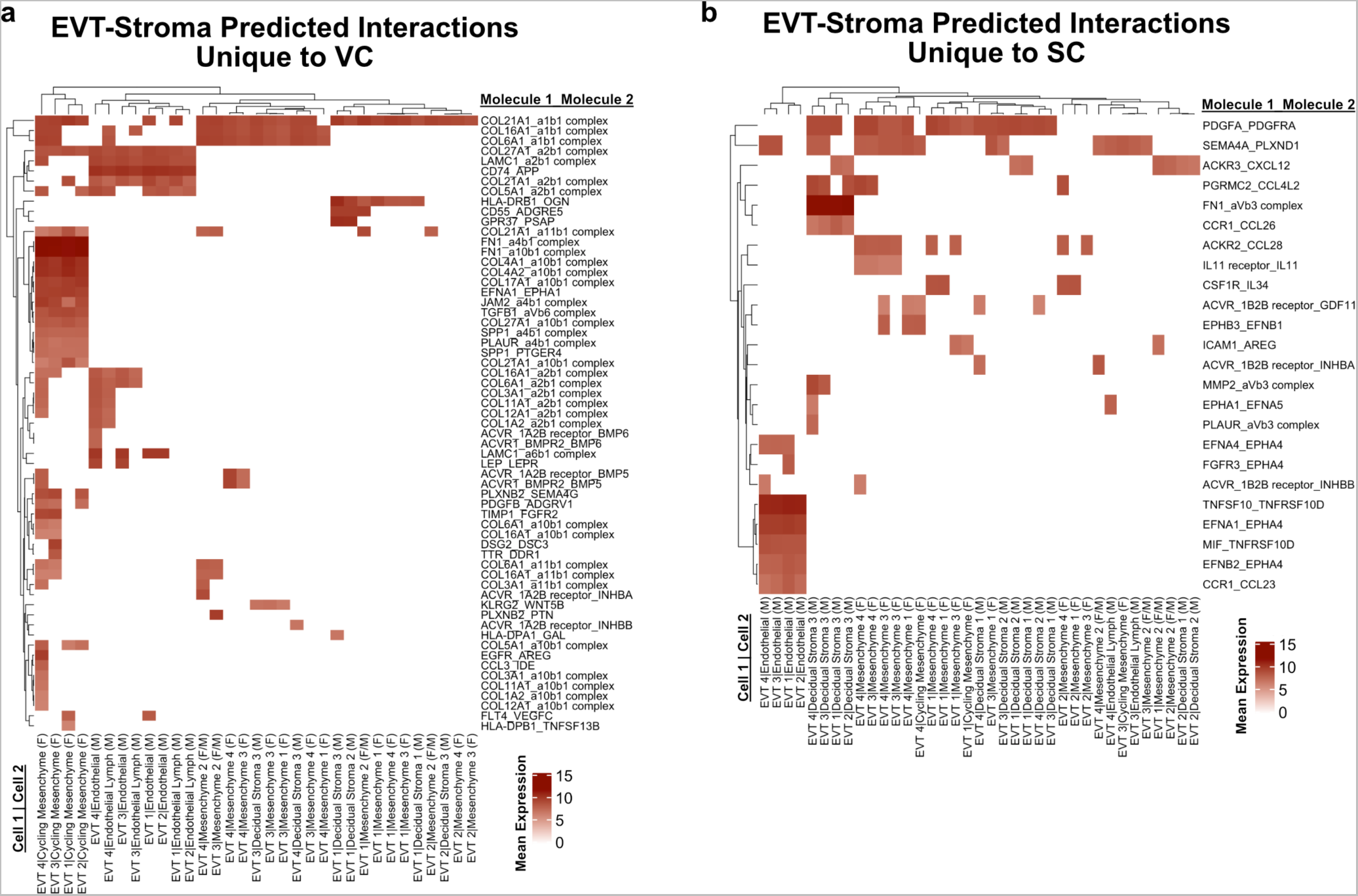
Predicted interactions between EVT and stromal cells. **(a)** Predicted receptor-ligand interactions from CellPhoneDB between EVT-Stromal cells of the VC for interactions which are unique to the VC. **(b)** Predicted receptor-ligand interactions from CellPhoneDB between EVT-Stromal cells of the SC for interactions which are unique to the SC. The strength of interaction is estimated by mean expression and are plotted in the heatmaps. Receptor-ligand interactions and cell pairs are listed such that Molecule 1 is expressed by Cell 1 and Molecule 2 is expressed by Cell 2.

**Figure 6-S1.**
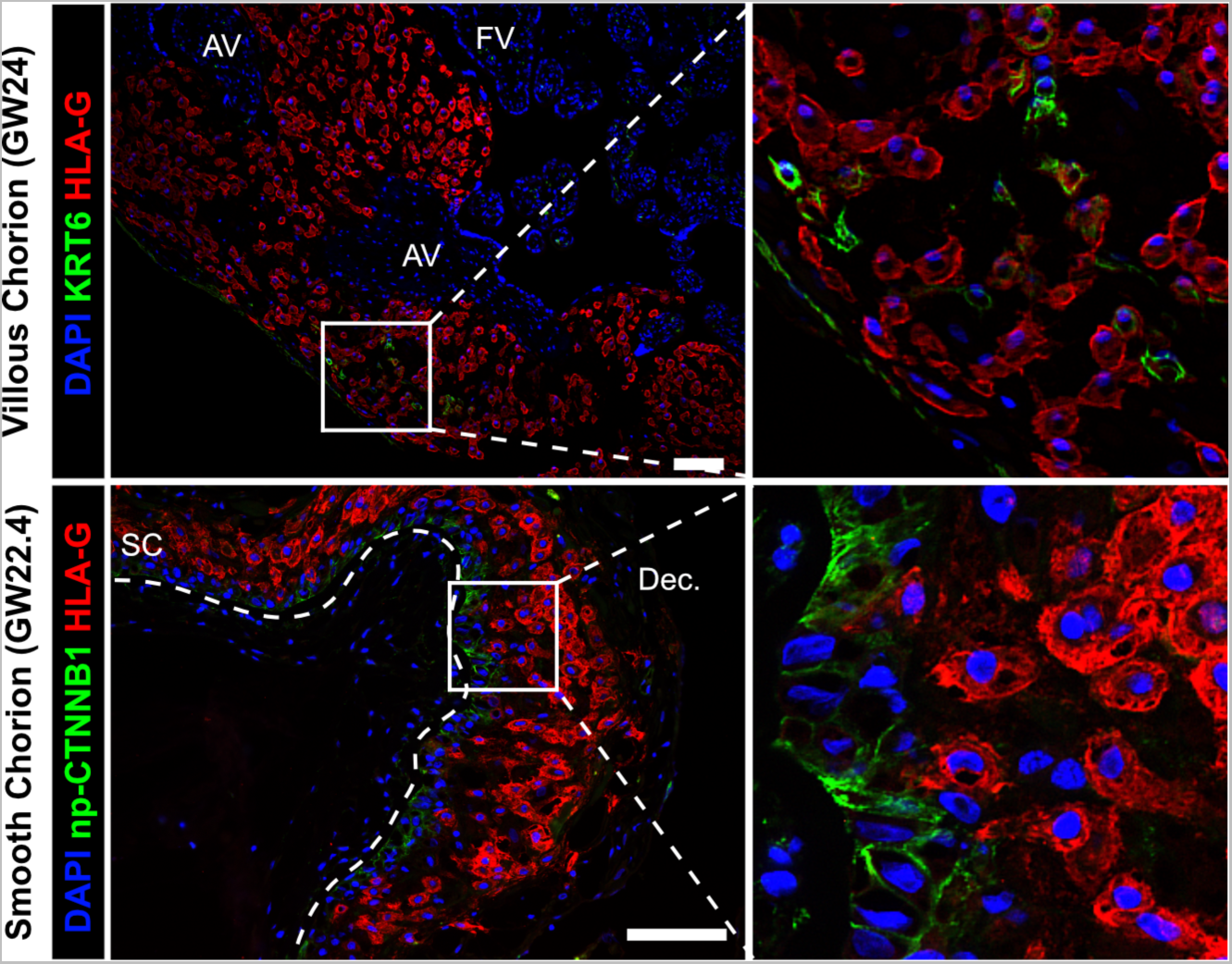
CTB-EVT interactions in the VC or SC region. Colocalization of KRT6 (SC-CTB marker) and HLA-G (EVT marker) showing rare KRT6+ cells in the VC (top). These cells are few and do not interact with EVT in the same manner as was observed in the SC. Co-localization of non-phosphorylated CTNNB1 (CTB 1 marker) and HLA-G (EVT marker) in the SC (Bottom). Limited interactions between these populations was observed. The basal lamina separating the fetal stroma from the SC epithelium is marked by the white dashed line. For all images nuclei were visualized by DAPI stain; Scale bar = 100μm. Abbreviations: AV = Anchoring Villi; FV = Floating Villi; SC = Smooth Chorion epithelium; Dec. = Decidua.

**Figure 6-S2.**
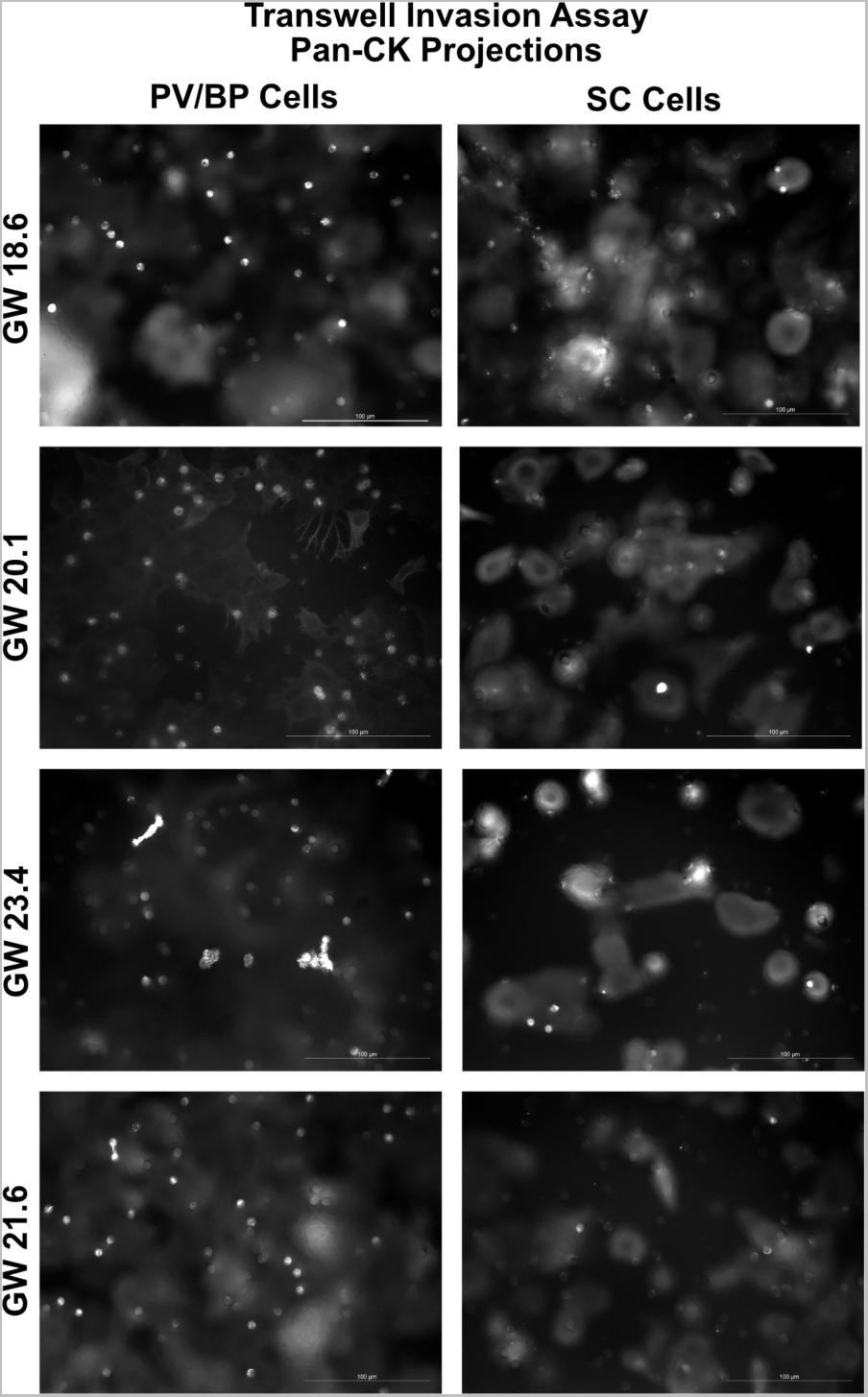
Representative images of the transwell invasion assay. Representative images of trophoblast projections through the transwell filter are seen a white dots. For quantification, the number of projections (white bright dots) were counted. For each sample, the median DAPI area was quantified across the transwell membrane, and then the projections multiplied by a factor to normalize for cell density. Scale bar = 100μm.

**Figure 6-S3.**
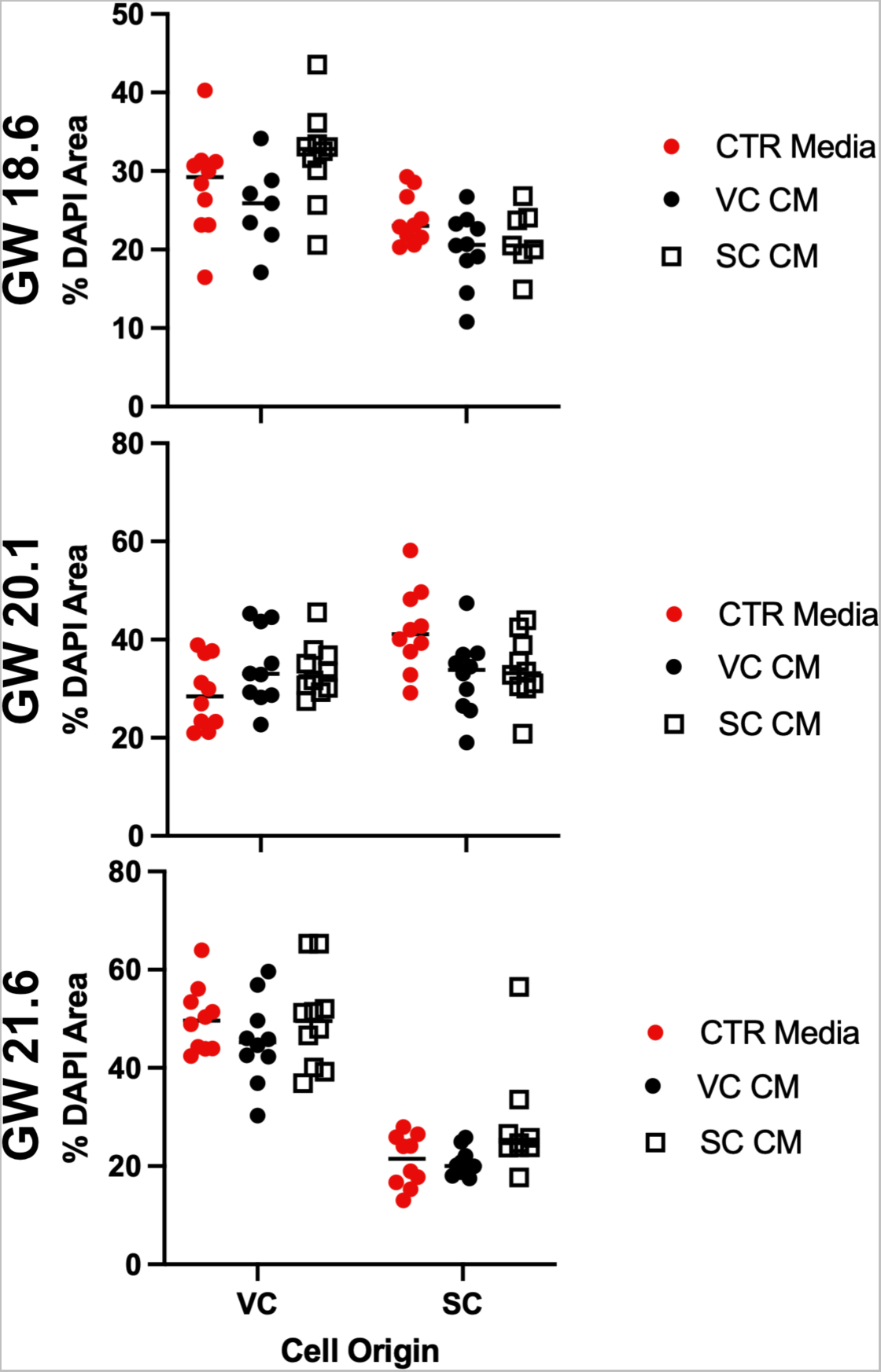
Cell density is not correlated with culture in conditioned media. The DAPI+ area in each field of view was quantified and plotted as a measure of cell density. Each dot represents the percent area of the field of view which stained with DAPI. Measurements from each experiment are shown and the gestational ages of the cultured cells is on the y-axis at the left. Measurements of cells treated with control media (Serum Free Media) are shown in red, VC cell conditioned media are shown in black, and SC cell conditioned media are shown in open squares.

**Figure 6-S4.**
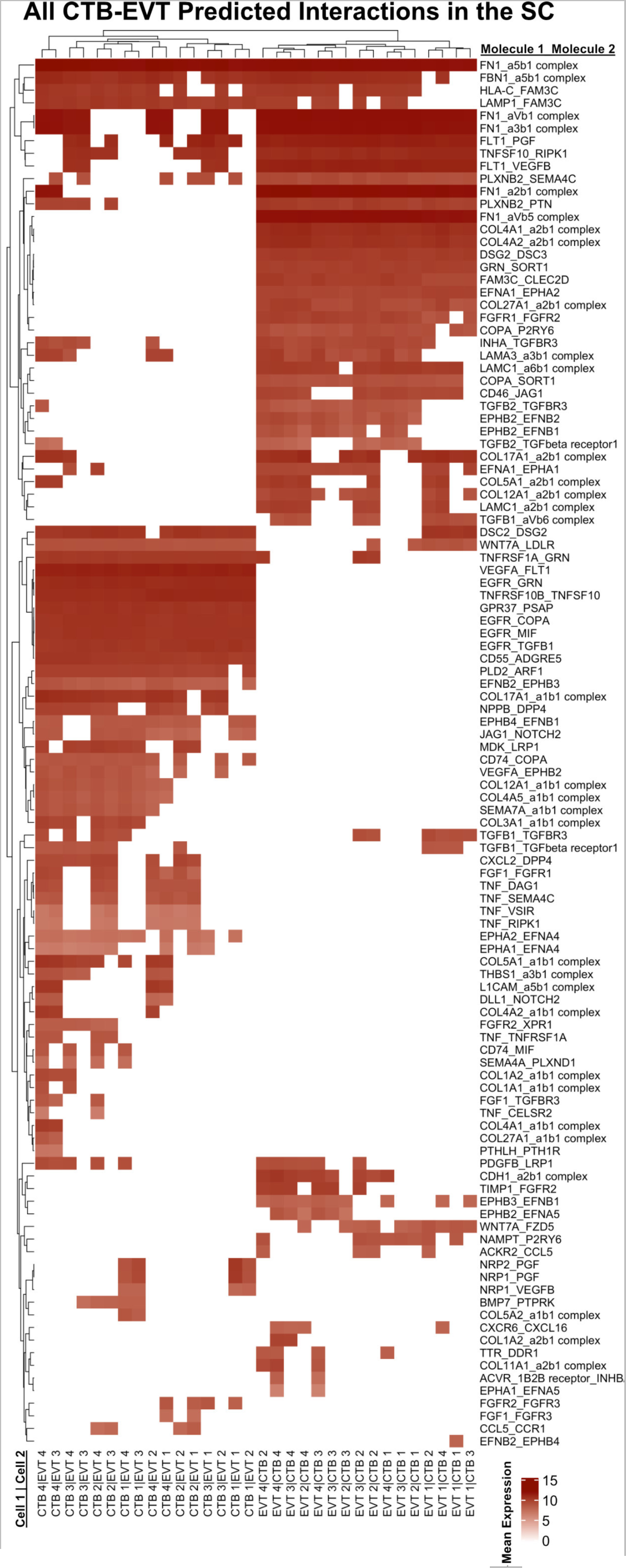
Complete predicted interactions between CTB-EVT in the SC. **(a)** Predicted receptor-ligand interactions from CellPhoneDB between CTB-EVT the SC. The strength of interaction is estimated by mean expression and are plotted in the heatmaps. Receptor-ligand interactions and cell pairs are listed such that Molecule 1 is expressed by Cell 1 and Molecule 2 is expressed by Cell 2.

